# Bayesian variable selection with a pleiotropic loss function in Mendelian randomization

**DOI:** 10.1101/593863

**Authors:** Apostolos Gkatzionis, Stephen Burgess, David V Conti, Paul J Newcombe

## Abstract

Mendelian randomization is the use of genetic variants as instruments to assess the existence of a causal relationship between a risk factor and an outcome. A Mendelian randomization analysis requires a set of genetic variants that are strongly associated with the risk factor and only associated with the outcome through their effect on the risk factor. We describe a novel variable selection algorithm for Mendelian randomization that can identify sets of genetic variants which are suitable in both these respects. Our algorithm is applicable in the context of two-sample summary-data Mendelian randomization and employs a recently proposed theoretical extension of the traditional Bayesian statistics framework, including a loss function to penalize genetic variants that exhibit pleiotropic effects. The algorithm offers robust inference through the use of model averaging, as we illustrate by running it on a range of simulation scenarios and comparing it against established pleiotropy-robust Mendelian randomization methods. In a real data application, we study the effect of systolic and diastolic blood pressure on the risk of suffering from coronary heart disease. Based on a recent large-scale GWAS for blood pressure, we use 395 genetic variants for systolic and 391 variants for diastolic blood pressure. Both traits are shown to have significant risk-increasing effects on coronary heart disease risk.

## 1 Introduction

Mendelian randomisation provides a framework for probing questions of causality from observational data using genetic variants. It applies the theory of instrumental variable analysis from the causal inference literature, using genetic variants associated with the risk factor as instruments. Mendelian randomization relies on the idea that, since genetic variants are randomly inherited and fixed at conception, they should be uncorrelated with potential confounders of the relationship between the risk factor and outcome and are therefore suitable to use as instruments. This approach has received much attention since the seminal paper of Davey Smith and Ebrahim (2003), and has led to a number of influential results over the last decade addressing a variety of aetiological questions (Boef et al., 2015). For example, in coronary heart disease, Mendelian randomization has been used to strengthen the evidence for a causal role of lipoprotein(a) (Kamstrup et al., 2009), but to weaken the case for C-reactive protein (CRP CHD Genetics Collaboration, 2011).

Formally, Mendelian randomization relies on three basic assumptions:

1. There is no confounding in the association between the genetic variants and the outcome.
2. The genetic variants are associated with the risk factor of interest.
3. The genetic variants only influence the outcome via their association with the risk factor and not through alternative causal mechanisms.

The three assumptions are illustrated in Figure 1. Under these assumptions, valid causal inferences can be made as to whether the risk factor affects the outcome.

**Figure 1:**
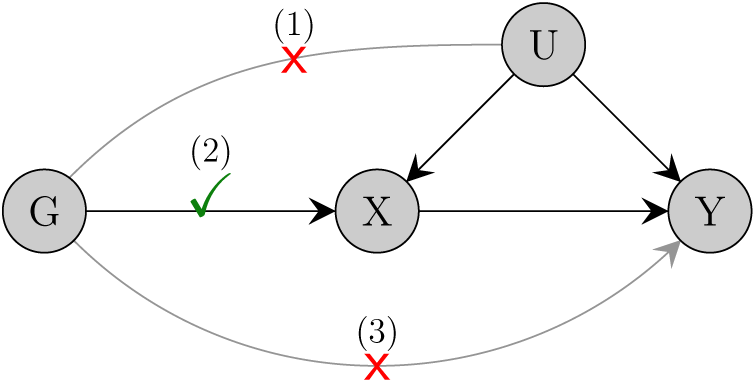
A causal diagram representation of the three assumptions of Mendelian randomization. Here, *X* represents the risk factor, *Y* the outcome, *G* the genetic instrument and *U* denotes confounders of the *X* − *Y* relationship.

As mentioned earlier, assumption (1) is usually justified on the basis of Mendelian inheritance: an individual’s genotype is randomly assorted at conception and not influenced by external confounding factors. To ensure the validity of assumption (2) Mendelian randomization analyses utilize the results of large consortia meta-GWAS, which use sample sizes of tens or hundreds of thousands of individuals to identify genetic variants robustly associated with a trait. In particular, many recent Mendelian randomization studies have taken a two-sample approach in which genetic associations with the target risk factor and with the outcome are assessed in separate datasets, in order to leverage their respective (and often mutually exclusive) meta-GWAS. However, these GWAS results are rarely available as individual-level data. Usually, only a set of summary statistics, such as univariate SNP-trait associations and corresponding standard errors, are reported. As a result, a large number of recent Mendelian randomization investigations rely on summarized data.

Assumption (3), often called the exclusion restriction or no-pleiotropy assumption, has received much attention in the recent literature. In practical applications, we typically do not know which genetic variants exhibit pleiotropic effects and there is need for methods to perform Mendelian randomization in the presence of pleiotropic variants. Traditional approaches include MR-Egger regression (Bowden et al., 2015) and median estimation (Bowden et al., 2016), while several algorithms for pleiotropy-robust Mendelian randomization have been developed recently (Hartwig et al., 2017; Kang et al., 2016; Rees et al., 2019; Burgess et al., 2018; Verbanck et al., 2018; Zhao et al., 2018; Qi and Chatterjee, 2018; Bowden et al., 2018; Burgess et al., 2019). Recent reviews and comparison of the various methods can be found in Slob and Burgess (2019) and Qi and Chatterjee (2019).

In this paper, we add to the relevant literature by proposing a new method for variable selection and causal effect estimation in Mendelian randomization. Our method is derived as an extension of the JAM algorithm (Joint Analysis of Marginal summary statistics, Newcombe et al., 2016). JAM was originally proposed for fine-mapping genetic regions. Similar to other recently proposed fine-mapping algorithms (Benner et al., 2016; Chen et al., 2015), JAM is designed to work with summary GWAS data. The algorithm performs variable selection to identify genetic variants robustly associated with the trait. Genetic correlations are taken into account by estimating them from a reference dataset such as 1000 Genomes or the UK Biobank. Variable selection is performed according to a Bayesian stochastic search algorithm, which can explore the complete space of causal configurations. Consequently, JAM is able to explore complex models with large numbers of variants, as recently demonstrated while fine-mapping dense genotype data for prostate cancer risk (Dadaev et al., 2018).

We develop a novel model averaging variable selection algorithm for Mendelian randomization, which we call JAM-MR (JAM for Mendelian Randomization). To do so, we modify JAM’s variable selection to downweight genetic variants which exhibit heterogeneous effects. We use the recently proposed framework of general Bayesian inference (Bissiri et al., 2016), in which the likelihood is augmented with a loss function before obtaining a Bayesian loss-posterior distribution. JAM’s likelihood is hence combined with a loss function that penalizes models containing variants with heterogeneous univariate causal effect estimates. Our algorithm performs variable selection and returns variant-specific posterior inclusion probabilities, which can be interpreted as probabilities of each variant being a valid instrument, and posterior model probabilities, which can subsequently be used to estimate the causal effect of interest by averaging across model-specific estimates. Uncertainty in which variants should be excluded on the basis of pleiotropy is reflected by averaging estimates over competing selections of instruments; model averaged causal inference is an attractive feature and one of the key contributions of our method.

Our algorithm has a number of additional advantages compared to established Mendelian randomization approaches. The Bayesian stochastic search used by JAM-MR has already proven useful in other areas of statistical genetics, such as fine-mapping (Benner et al., 2016; Newcombe et al., 2016) and colocalization (Asimit et al., 2019). The use of the Bayesian paradigm allows us to incorporate prior information on the suitability of genetic variants as instruments into the analysis. It also allows us to model the uncertainty in genetic associations with the risk factor, which is often ignored by other approaches. JAM-MR also offers a natural framework for incorporating genetic correlations, when conducting Mendelian randomization with several genetic variants coming from the same gene region.

The performance of our algorithm is illustrated in a range of simulated datasets, as well as in a real data application where we investigate the causal effect of systolic and diastolic blood pressure on the risk of coronary heart disease. We instrument blood pressure using a recently published large scale meta-GWAS, which combined results across more than one million individuals, and therefore base our Mendelian randomization analysis on larger sample sizes and more genetic variants compared to previous studies in the literature. Our results strengthen the claim for a risk-increasing causal relationship between blood pressure traits and coronary heart disease risk.

## 2 Methods

### 2.1 The JAM algorithm

#### 2.1.1 Introduction

We start by providing a brief outline of the JAM algorithm. The reader is referred to Newcombe et al. (2016) for a more detailed description.

The JAM algorithm is primarily a tool for fine-mapping densely genotyped DNA regions. The standard fine-mapping question is as follows: given *P* correlated genetic variants *G*_1_, …, *G*_*P*_, identify which variants are causally and independently associated with a trait *X* of interest. The genetic variants are typically located in the same gene region. The main purpose of JAM is to perform variable selection for the genetic variants, given a set of GWAS summary statistics and an external dataset from which to estimate genetic correlations.

For individual *i*, let *g*_*i*_ = (*g*_*i*1_, …, *g*_*iP*_) and *x*_*i*_ be the allele counts and trait measurements respectively, and denote *G* = (*g*_*ij*_) the genetic matrix. We assume that trait values and allele counts per variant have been centered. Typically, the “individual-level data” *x*_*i*_, *g*_*i*_ are not available in practice. Instead, we only have access to a set of univariate association estimates 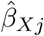 between each variant and the trait, as well as the corresponding standard errors 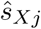.

#### 2.1.2 The JAM model

JAM uses linear regression to model the trait: if all *P* genetic variants were assumed to be associated with *X*, the JAM algorithm would model the trait as

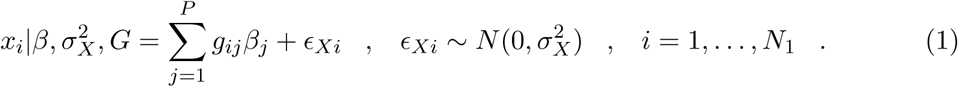

In practice JAM implements variable selection, reflecting that, in many applications, only a subset of the variants should be used to model the trait. Let *γ* ∈ {0, 1}^*P*^ index the selected subset, so that if *γ*_*j*_ = 1 variant *G*_*j*_ is included into the JAM model and if *γ*_*j*_ = 0 it is not. Using *β*_*γ*_, *G*_*γ*_ to denote the subsets of *β, G* only for the variants *G*_*j*_ for which *γ*_*j*_ = 1, (1) becomes

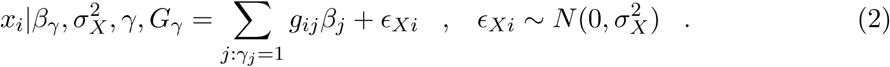

Equation (2) can be used to build a likelihood for the individual-level data,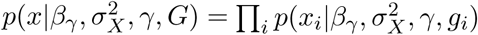. One can also obtain the marginal model likelihood,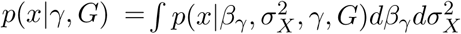. JAM works by constructing summary-data approximations to these two likelihoods (see the following subsections).

#### 2.1.3 Prior specification

The likelihood 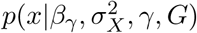 is complemented with a set of priors in order to perform Bayesian inference. For the genetic associations *β*_*γ*_, JAM uses a conjugate g-prior,

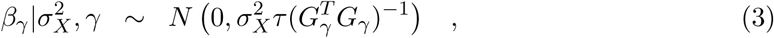

where *τ* is a constant. By default, the algorithm sets *τ* = max(*P*^2^, *N*_1_), as has been previously recommended by various authors (George and McCulloch, 1997; Fernandez et al., 2001). The residual variance 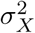 is assigned its own conjugate prior, which is an Inverse-Gamma density,

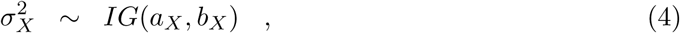

for fixed *a*_*X*_, *b*_*X*_. Finally, JAM uses a Beta-Binomial prior on the space of all possible models,

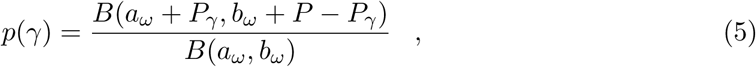

where *B*(*a, b*) denotes the Beta function and *P*_*γ*_ denotes the size of model *γ*. The relative sizes of the hyperparameters *a*_*ω*_, *b*_*ω*_ reflect the proportion of genetic variants expected a priori to be associated with the trait; by default, *a*_*ω*_ = 1, *b*_*ω*_ = *P*, which correspond to an expectation that a single variant will be associated with the trait.

#### 2.1.4 Posterior inference for the regression parameters

A Bayesian posterior distribution over all the parameters (model indicator, genetic associations and residual variance) can be obtained according to the standard principles of Bayesian inference:

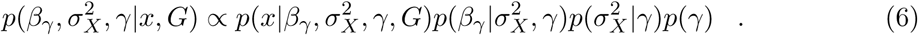

Conditional on a particular combination of causal variants, posterior inference on the regression parameters 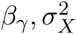 can be conducted using known results for Bayesian linear regression with conjugate priors, which yield

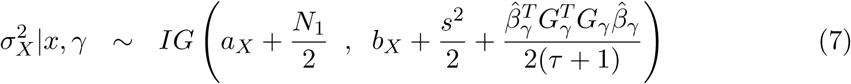

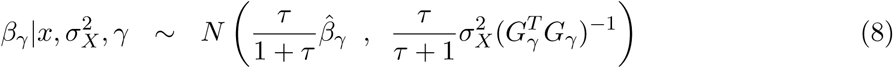

Where 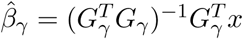 and 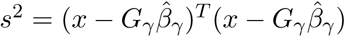.

#### 2.1.5 Posterior model selection

An advantage of the linear regression setting is that it allows for fast and efficient variable selection. The regression coefficients *β*_*γ*_ and the residual variance 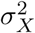 can be integrated out from the JAM likelihood 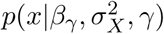 to obtain the marginal model likelihood

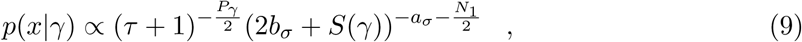

where

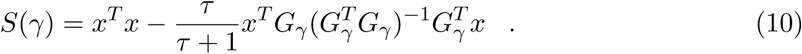

This leads to the marginal model posterior, *p*(*γ*|*x*) ∝ *p*(*x*|*γ*)*p*(*γ*). The normalizing constant of that density can be difficult to evaluate, but this can be avoided using reversible-jump MCMC (Green, 1995). JAM implements a standard reversible-jump algorithm with addition, deletion and swapping of genetic variants as possible moves. The stochastic search algorithm allows exploration of an unrestricted model space, without the need to set limits on the maximum number of causal variants. Consequently, JAM is able to efficiently explore complex causal configurations among large numbers of genetic variants. Posterior model probabilities can be estimated by the proportion of iterations JAM spends in each model.

#### 2.1.6 JAM with summarized data

One of JAM’s main advantages is that it does not require access to individual-level data. Variable selection is implemented according to Equations (9), (10), which depend on the observed data *x*_*i*_, *g*_*i*_ only through the quantities *x*^*T*^ *x, G*^*T*^ *x* and *G*^*T*^ *G*. These quantities are sufficient summary statistics for linear regression. In the setting of genome-wide association studies, Newcombe et al. (2016) describes a way of approximating *z* = *G*^*T*^ *x* from the univariate variant-trait association estimates 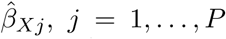, as well as effect allele frequencies. In addition, note that 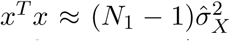 since we have assumed that trait values have been centered before implementing JAM. Here,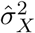 is an estimator of the trait variance Var(*X*) measured in the GWAS for the trait *X*. This variance is sometimes reported by genetic association studies, but can be approximated using the SNP-trait associations, standard errors and effect allele frequencies if it is not available (Yang et al., 2012). Finally, the matrix *G*^*T*^ *G* models (*N*_1_ − 1 times) the genetic correlations between variants *G*_1_, …, *G*_*P*_ and can be approximated if a reference dataset, such as the 1000 Genomes dataset or the UK Biobank, is available. This yields the following summary-data approximation for *S*(*γ*):

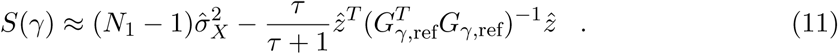

The model-specific marginal posteriors (7) and (8) can be approximated by summary GWAS data in a similar way.

#### 2.1.7 Binary traits

The JAM algorithm assumes that the trait of interest is continuous, as is often the case for risk factors used in Mendelian randomization studies. JAM then relies on a linear modelling framework to relate univariate summary statistics to multivariate effects via the transformation *z* = *G*^*T*^ *x*. If the trait studied is binary, *G*^*T*^ *x* can be derived by mapping univariate log-odds ratios to the univariate effects that would have been estimated if the binary trait was modelled by linear regression. The same strategy is employed in other linear-based summary data frameworks, including LDPred (Vilhjalmsson et al., 2015), to which we refer readers for a detailed description of this mapping.

### 2.2 An extension of JAM for Mendelian randomization

#### 2.2.1 Scope

We now describe our extension to the standard JAM algorithm that facilitates pleiotropy adjustment and accurate causal effect estimation in Mendelian randomization studies, which we call JAM-MR (JAM for Mendelian Randomization).

We start by pointing out a difference in the scope of our new algorithm compared to the original JAM algorithm. Traditionally, JAM is used to analyze correlated variants from one (fine-mapping) or multiple genetic regions. On the other hand, Mendelian randomization studies typically use variants from across the whole genome, often pruned for independence. Consequently, many practical implementations of JAM-MR will rely on independent, genome-wide significant variants. Mendelian randomization analyses using correlated variants are less common in practice, and assessing pleiotropy in these studies can be difficult because many genetic variants may share a common pleiotropic effect through correlation. Although JAM-MR is applicable in such studies too, we mainly consider the “traditional” Mendelian randomization framework in this paper.

For the JAM-MR algorithm, we work in the context of two-sample summary-data Mendelian randomization. This assumes that we have access to two different genetic association studies, one from which to obtain the SNP-risk factor summary statistics 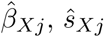 and another study from which to obtain the SNP-outcome univariate association estimates 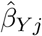 and standard errors 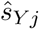. This two-sample summary-statistics framework is a common approach for Mendelian randomization, since it allows researchers to leverage the power and large sample sizes of large consortia GWAS studies that are already available for many important traits, without the requirement that both the risk factor and the outcome should be measured on the same GWAS.

The coefficients 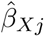 and 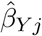 can be used to obtain variant-specific estimates of the causal effect *θ* according to a ratio formula (Burgess et al., 2017),

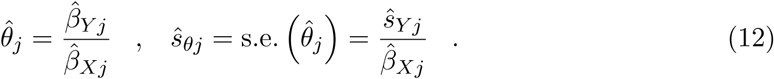

#### 2.2.2 Bayesian inference with loss functions

The JAM-MR algorithm implements a recently proposed framework for performing Bayesian inference using loss functions (Bissiri et al., 2016). This new theoretical framework constitutes a generalization of the core Bayesian paradigm. For a dataset *𝒟*, parameter vector *θ* and prior distribution *π*(*θ*), the standard Bayesian updating scheme, *p*(*θ*|*𝒟*) ∝ *π*(*θ*)*p*(*𝒟*|*θ*), is replaced with a loss-function update of the form

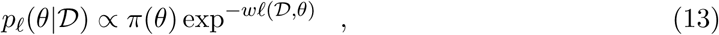

where *ℓ*(*𝒟*|*θ*) represents a loss function. Here, *w* is a tuning parameter that determines the influence of the loss function relative to that of the prior. If *w* = 1 and *ℓ*(*𝒟, θ*) = − log *p*(*𝒟*|*θ*) is the negative log-likelihood, we obtain traditional Bayesian inference.

The motivation behind this approach is that the loss function can be used to tailor Bayesian inference towards specific objectives. For example, if the objective of interest is classification, the misclassification error can be used as a loss function (Jiang and Tanner, 2008). In addition, using a loss function instead of a likelihood avoids the need for the Bayesian statistician to specify a full data-generating model; this is especially useful when the object of interest is a low-dimensional parameter within a complex, high-dimensional model. It also provides robustness to model misspecification, since the statistician is required to make fewer modelling assumptions. A rigorous decision-theoretic framework for general Bayesian inference is provided in Bissiri et al. (2016).

#### 2.2.3 JAM with a pleiotropic loss function

In order to construct the JAM-MR algorithm, we use a slight modification of the Bayesian loss function framework. Specifically, we use both a likelihood and a loss function to construct the “loss-posterior”:

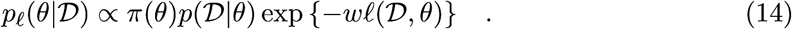

This is equivalent to using 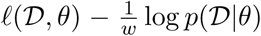 as a loss function in (13). The use of both a likelihood and a loss function in JAM-MR is justified because the objective of the model selection procedure is two-fold: we use the likelihood to select genetic variants strongly associated with the risk factor and the loss function to penalize variants which exhibit pleiotropic effects on the outcome. The parameter *w* can be interpreted as a tuning parameter that balances the impact of the pleiotropic loss on model selection relative to that of the JAM likelihood and the prior. Note that the loss function framework allows us to avoid making specific modelling assumptions for the SNP-outcome association, since that association is only modelled through the loss function.

In the context of JAM-MR, the data to be used are the univariate summary statistics along with the reference dataset from which to compute genetic correlations, so 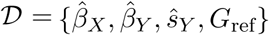. For the purpose of model selection, and since *β*_*γ*_ and 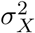 can be integrated out, the parameter vector is simply the model indicator *θ* = *γ* and the “likelihood” to be used in (14) is the marginal model likelihood (9); technically this is not a full likelihood, as it only models the *G* − *X* associations and not the *X* − *Y* causal relationship.

We now discuss how to specify the loss function *ℓ*(*𝒟, γ*). It is common in the Mendelian randomization literature to use some measure of heterogeneity between univariate causal effect estimates as a proxy for pleiotropic behaviour (Hartwig et al., 2017; Burgess et al., 2018). The intuition is that genetic variants which are valid instruments yield the same univariate causal effect estimates, up to some random variation. On the other hand, estimates based on pleiotropic variants can exhibit systematic differences, especially if the variants operate on different causal pathways towards the outcome, because the estimated causal effects depend on the strength and direction of the pleiotropic G-Y association. This suggests that our loss function should upweight models with homogeneous univariate causal effect estimates, as such models are likely to contain valid instruments, and downweight models with heterogeneous estimates, as at least some of the genetic variants contained in them are likely to be pleiotropic.

Consequently, in order to penalize pleiotropic models, we need a loss function that measures the heterogeneity of univariate estimates. A simple option is the sum of squared differences,

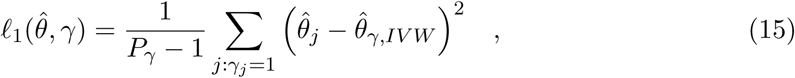

where

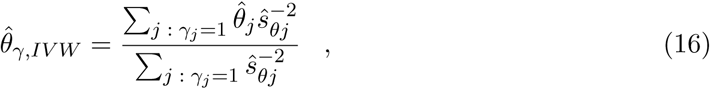

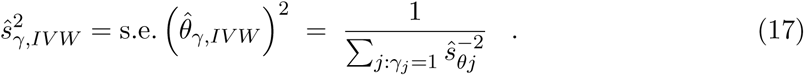

is the inverse-variance-weighted (IVW) causal effect estimate based on variants in model *γ*. The loss function downweights models with heterogeneous causal effect estimates, since the sum-of-squares in (15) is larger and the term exp{−*wℓ*(*𝒟, θ*)} in (14) is smaller. Note that the loss function is not defined for the null model and models containing only one genetic variant. Such models carry no evidence as to whether the variant included is valid or pleiotropic. Therefore, we ignore these models and restrict JAM-MR to only consider models with at least two genetic variants.

An alternative loss function can be obtained by weighting the individual causal effect estimates in (15) by the inverse of their squared standard errors,

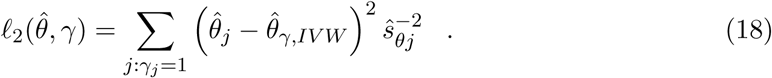

This second loss function has the advantage that it also models uncertainty in the univariate ratio estimates 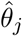 through its dependence on the standard errors 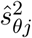. A similar function was used in Burgess et al. (2018) to obtain a pleiotropy-robust exhaustive-search model averaging procedure for Mendelian randomization. In the simulations and the real-data application of this paper, we have used the weighted loss function (18).

In conclusion, the Bayesian loss-posterior for JAM-MR’s variable selection is

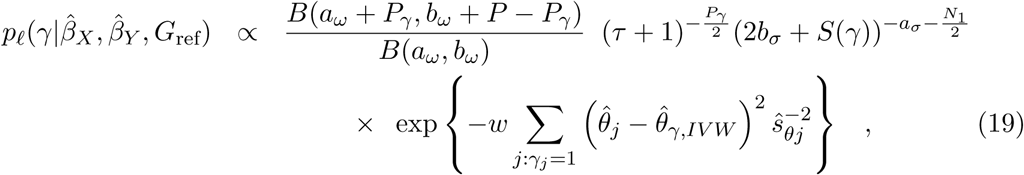

where *S*(*γ*) can be computed from (11). Note that this formula still depends on the tuning parameter *w*; the specification of a value for *w* will be discussed in Section 2.2.5. Given a fixed value for *w*, the loss-posterior (19) is used as the target distribution for JAM-MR’s reversible-jump MCMC algorithm. The stochastic search procedure scales well to large datasets containing hundreds of genetic variants, is quite flexible and can explore large parts of the model space.

Since we are using heterogeneity as a proxy for pleiotropic behaviour, JAM-MR implicitly makes a plurality assumption: the algorithm targets the largest set of genetic variants with homogeneous univariate causal effect estimates, and it is assumed that this set corresponds to the valid instruments. This assumption is similar to those made in Hartwig et al. (2017) and Burgess et al. (2018). If smaller sets of genetic variants with homogeneous ratio estimates exist, the stochastic search will sometimes identify them; in practice, this can be checked by inspecting the list of posterior model probabilities. Such sets may correspond to several pleiotropic variants acting on the same causal pathway to the outcome, and the biological interpretation of these sets can be an interesting question in applications. We note however that JAM-MR often requires a very large number of iterations in order to identify such sets.

#### 2.2.4 Causal effect estimation using JAM-MR

The JAM-MR posterior probabilities can subsequently be used to obtain an overall estimate of the causal effect of interest, according to a model averaging procedure. The use of Bayesian model averaging enables quantification of uncertainty related to the choice of instruments and allows many genetic variants to have small contributions to the overall causal effect estimate.

For each model *γ*, the algorithm computes a model-specific causal effect estimate 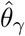 and standard error *ŝ*_*γ*_. These are then combined into a single estimator,

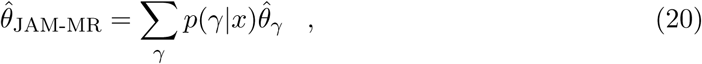

with variance

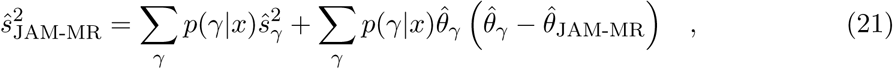

where the summation is over all models *γ* assigned positive posterior probability by the variable selection procedure. Equation (21) can be derived as an approximation to the posterior variance Var(*θ*|*𝒟*), by expressing it in terms of the model-specific posterior moments 𝔼(*θ*|*𝒟, γ*), Var(*θ*|*𝒟, γ*) and approximating these by 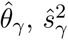.

A straightforward choice for the model-specific estimates would be the inverse-variance-weighted estimate (16). However, we have observed empirically that this choice often results in underestimation of the overall causal standard error and a more elaborate modelling assumption may be needed. To motivate that, consider the simulation plotted in Figure 2. The figure plots univariate causal effect estimates and standard errors for *P* = 50 genetic variants. Variants 1-35 were generated as valid instruments and variants 36-50 were generated to be pleiotropic. The true causal effect of the risk factor on the outcome was set to zero. JAM-MR was implemented to identify variants suitable for inclusion in a Mendelian randomization analysis. The variants that were assigned a posterior inclusion probability lower than 0.5 are coloured red.

**Figure 2:**
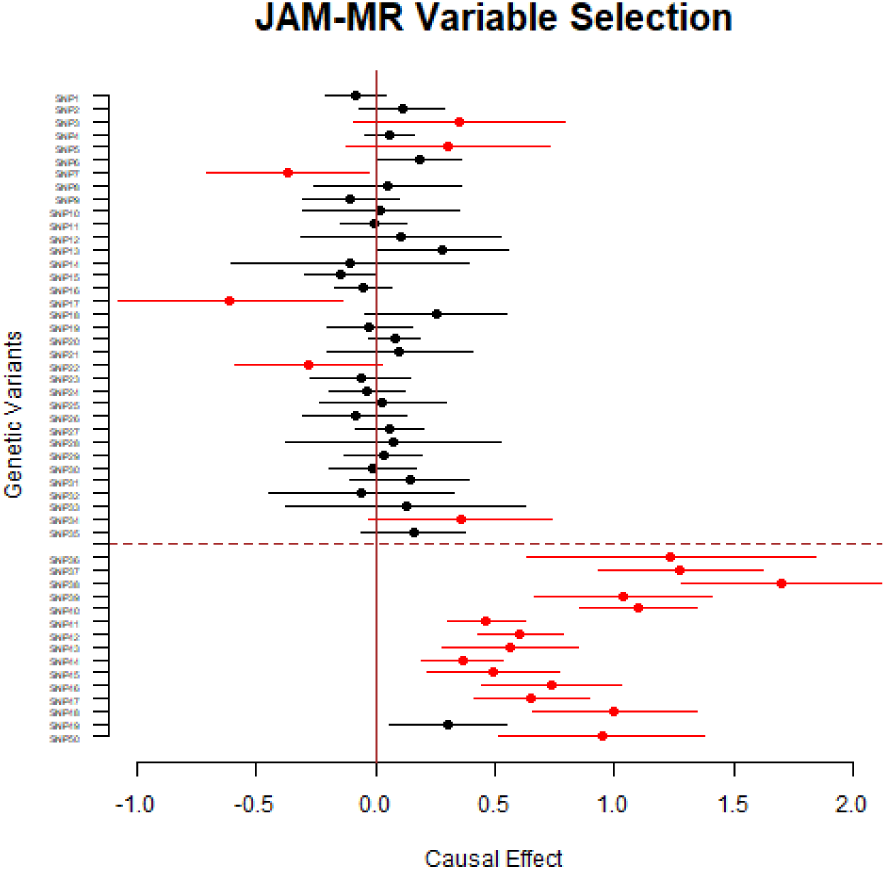
JAM-MR variable selection in a simulated dataset of 50 genetic variants. A single implementation of a directional pleiotropy simulation (scenario 2) with *θ* = 0. Causal effect estimates and 95% confidence intervals for each variant are plotted. Variants above the dotted line were simulated as valid and variants below the dotted line were simulated as pleiotropic. Variants assigned a posterior inclusion probability lower than 50% are coloured red.

Because JAM-MR uses the heterogeneity of ratio estimates in order to model pleiotropic behaviour, it will tend to favour sets of variants with homogeneous causal effect estimates and consequently under-estimate the overall heterogeneity. This manifests as overly-precise standard errors, as estimated from the naive IVW formula. To better model this behaviour and attenuate the overprecision, we modify the inverse-variance-weighted estimates as follows.

The standard IVW formula can be motivated as the maximum likelihood estimate obtained from the heteroskedastic linear regression model 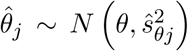. To account for the presence of moderately pleiotropic variants, which may have been inadvertently included in JAM-MR’s model due to their proximity to valid instruments, it has been proposed Burgess and Bowden (2016) to use a multiplicative random effects model of the form 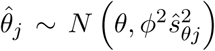. This results in the same causal effect estimate (16) as the inverse-variance-weighted approach, but scales its standard error by *ϕ*. The overdispersion parameter *ϕ* can be estimated using weighted linear regression (Burgess and Bowden, 2016). On the other hand, to model the potential downweighting of valid genetic variants with outlying ratio estimates, we modify the random-effects model further and use a truncated normal distribution for the univariate causal effects:

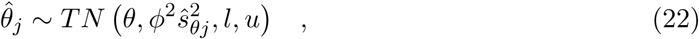

where *l, u* denote the lower and upper truncation points. Using (22) we can obtain a model-specific likelihood

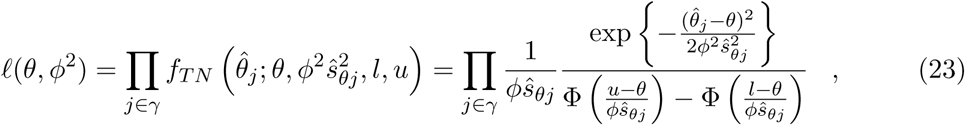

where Φ denotes the cumulative distribution function of a *N* (0, 1) random variable. The likelihood (23) can then be maximized numerically to obtain a model-specific estimate 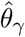 of the causal effect. To simplify the optimization problem, we fix the left truncation point *l* to the value 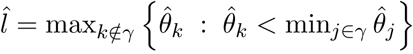. This is the equivalent of truncating at the largest ratio estimate 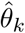 of a SNP not in model *γ* that is smaller than all the ratio estimates of SNPs in *γ*. If no such 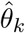 exists, we do not truncate to the left 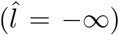. Likewise, we fix the right truncation point at 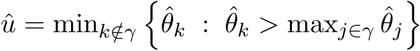, the smallest ratio estimate of a SNP not in model *γ* that is still larger than the ratio estimates of all SNPs in *γ* (or *û* = ∞, if no such 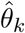 exists).

Optimization of (23) over (*θ, ϕ*^2^) can be performed using gradient-based methods, since the derivatives of the likelihood (23) are straightforward to compute. To estimate the causal standard error, we rely on frequentist asymptotic arguments and compute the sample Fisher information matrix,

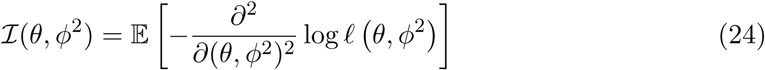

evaluated at the maximum likelihood estimates 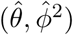. The maximum likelihood estimate for *θ* and its standard error are then used as the model-specific estimates in (20), (21), from which to obtain the overall causal effect estimate.

In the simulations and the real-data application of this paper, we have used the truncated multiplicative random-effects model in order to compute model-specific causal effect estimates and standard errors. A comparison between the three approaches (fixed-effects IVW, random-effects IVW and the truncated model) is provided in the Appendix.

Regardless of the modelling assumptions used, it is worth noting that the algorithm is not fully Bayesian. Although JAM-MR utilizes Bayesian variable selection to obtain posterior model probabilities and facilitate model averaging, the process of causal effect estimation for each model is conducted using classical inverse-variance-weighted formulas. A fully Bayesian algorithm for Mendelian randomization would require further modelling assumptions for the causal effect parameter and return a posterior sample for that parameter. This is an interesting potential extension of JAM-MR, but is beyond the scope of this paper.

#### 2.2.5 Estimation of the tuning parameter *w*

The tuning parameter *w* plays a crucial role in JAM-MR’s variable selection. Tuning *w* is subject to a bias-variance trade-off. For relatively small values, the pleiotropic loss function has limited effect on the variable selection procedure and JAM-MR tends to favour larger models. These models may still include some pleiotropic variants, and the resulting causal effect estimates may exhibit bias. On the other hand, with a large value of *w* the algorithm favours models that contain no pleiotropic variants but may also ignore some of the valid instruments. In this case JAM-MR yields unbiased causal effect estimates, but these estimates may have large standard errors.

Setting *w* = 0 is equivalent to the standard JAM algorithm with no pleiotropy adjustment; the algorithm is similar to a simple IVW estimator except it downweights genetic variants that are weakly associated with the risk factor. The other extreme case is to assign a very large value to *w*. In this case the algorithm becomes very selective and prioritizes models containing only two genetic variants.

In order to balance between these two extremes, JAM-MR implements a grid search to select the value of the tuning parameter. A range of candidate values is specified and the stochastic search is run for each value separately. Candidate values for the grid search should scale according to the sample size *N*_1_, since the value of the likelihood also scales according to *N*_1_. By default, JAM-MR uses a grid with 25 points, spaced equally on the logarithmic scale between *w*_1_ = 0.01*N*_1_ and *w*_25_ = 100*N*_1_ (that is, log *w*_1_, …, log *w*_25_ are equally spaced), plus an implementation for *w*_0_ = 0.

Among the candidate values for *w*, the algorithm then selects the one for which the smallest causal standard error (21) is obtained. Intuitively, the collection of genetic variants that attains the smallest standard error should correspond to the set of valid instruments, as both the inclusion of a pleiotropic variant and the removal of a valid SNP from that model will increase the causal standard error. In our simulations, we have found this criterion to offer a reasonable compromise in the bias-variance tradeoff that underpins the selection of a value for the tuning parameter.

## 3 Simulation Study

### 3.1 Simulation setting

We conducted a simulation study in order to assess the performance of the JAM-MR algorithm and compare it to established approaches for Mendelian randomization. The Mendelian randomization model that we used for the simulations was

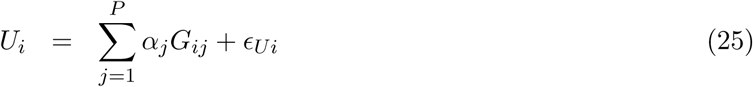

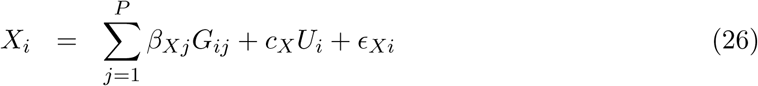

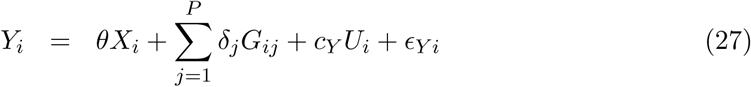

where 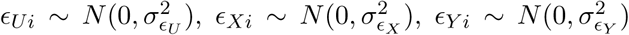 independently of each other. The effects of genetic variant *G*_*j*_ on *U, X, Y* are denoted by *α*_*j*_, *β*_*j*_, *δ*_*j*_ respectively. The parameters *δ*_*j*_ represent the direct effects of genetic instruments on the outcome. The parameters *α*_*j*_ represent effects mediated by confounders, which we use to model violations of the InSIDE assumption (“instrument strength is independent of direct effects”). For genetic variants that are valid instruments, *δ*_*j*_ = *α*_*j*_ = 0.

Our simulations were conducted using the statistical software R. To inform the simulations, we used sample sizes and numbers of genetic variants similar to those in the real data application presented in the next section. Specifically, we used a two-sample Mendelian randomization setting and generated two sets of individual-level data for *N*_1_ = *N*_2_ = 300000 individuals and *P* = 400 independent genetic variants. These sample sizes are in line with recent Mendelian randomization analyses using composite risk factors. Of the 400 genetic variants, *P*_1_ = 120 (30%) were simulated as pleiotropic and *P* − *P*_1_ = 280 were simulated as valid instruments.

Genetic variants *g*_*ij*_ were generated from a Binomial(2, *f*_*j*_), with effect allele frequencies *f*_*j*_ ∼ *U* (0.1, 0.9). The genetic effects on the risk factor were simulated as 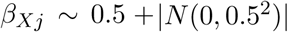, which we empirically observed to yield a realistic pattern of genome-wide significant p-values, the majority of which were between 10^−7^ and 10^−50^. The error term variances were set equal, 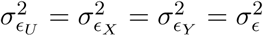, and the value of 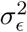 was specified so that 10% of the variation in the risk factor is explained by the genetic variants. We also set *c*_*X*_ = *c*_*Y*_ = 1.

The pleiotropic effect parameters *α*_*j*_, *δ*_*j*_ were set equal to zero for all the valid instruments. For invalid instruments, we generated three simulation scenarios. In the first scenario (balanced pleiotropy), we let *α*_*j*_ = 0 and *δ*_*j*_ ∼ ±*N* (0.7, 0.2^2^), setting the sign of each *δ*_*j*_ to be positive or negative with equal probability. In the second scenario (directional pleiotropy, InSIDE assumption satisfied) we also set *α*_*j*_ = 0 but the direct effect parameters *δ*_*j*_ were restricted to be positive, *δ*_*j*_ ∼ *N* (0.7, 0.2^2^). Finally, in the third simulation scenario (directional pleiotropy, InSIDE violated), the SNP-confounder effects were also randomly generated, *α*_*j*_ ∼ *N* (0.4, 0.2^2^). The direct effects *δ*_*j*_ were generated as in the second scenario. The three simulation scenarios are summarized in Table 1. Plots of univariate p-values for SNP-trait associations and univariate causal effect estimates for our simulations are provided in the Appendix.

**Table 1:** Different simulation scenarios.

| Scenario | Type of Pleiotropy | Direct Effects | Confounder Effects |
| --- | --- | --- | --- |
| 1 | Balanced | $\delta_j \sim \pm N(0.7, 0.2^2)$ | $\alpha_j = 0$ |
| 2 | Directional<br>InSIDE Satisfied | $\delta_j \sim N(0.7, 0.2^2)$ | $\alpha_j = 0$ |
| 3 | Directional<br>InSIDE Violated | $\delta_j \sim N(0.7, 0.2^2)$ | $\alpha_j \sim N(0.4, 0.2^2)$ |

For each scenario, we implemented two sets of simulations, one with a null (*θ* = 0) and one with a positive (*θ* = 0.3) causal effect. This resulted in a total of six simulation settings. Each setting was replicated 1000 times. In each replication, we generated the individual-level data and then performed univariate linear regression between each genetic variant and the risk factor and outcome to obtain GWAS summary statistics. Using these summary statistics, we then implemented JAM-MR as well as a range of already existing algorithms for Mendelian randomization. For comparison purposes, we also computed “oracle” IVW causal effect estimates using only the valid instruments. For more details on how JAM-MR and the competing methods were implemented in our simulation, we refer to the Appendix.

### 3.2 Competing methods

To assess the performance of the JAM-MR algorithm, we compared it against the following established methods for Mendelian randomization:

- Standard inverse variance weighted (IVW) estimation.
- MR-Egger regression (Bowden et al., 2015).
- Median estimation (Bowden et al., 2016).
- Mode-based estimation (Hartwig et al., 2017).
- A Lasso-type estimator (Rees et al., 2019).
- MR-Presso (Verbanck et al., 2018).
- MR-Raps (Zhao et al., 2018).
- Contamination mixture (Burgess et al., 2019).

A short description of each method is provided in the Appendix. For further details the reader is referred to the relevant citations.

### 3.3 Simulation results

The results of this simulation experiment are reported in Tables 2, 3 and 4 for each of the three simulation scenarios and either a null (*θ* = 0) or a positive (*θ* = 0.3) causal effect. We report average causal effect estimates, estimated standard errors and root mean squared errors. Standard errors for the contamination mixture are not reported because the method does not compute standard errors and directly obtains a confidence region (which is not necessarily a single interval) instead. In the case of a null causal effect, we also report Type I error rates (calculated as the empirical power to reject the null causal hypothesis) at a 95% significance level, while for simulations with a positive causal effect we report the empirical coverage of 95% confidence intervals. We do not report the power to reject the null hypothesis when it is not true, because all methods were able to do so in almost all the simulation replications; this was due to the large sample sizes that we used, in combination with *θ* = 0.3 being a rather strong effect that most methods could easily identify.

**Table 2:** Scenario 1: Balanced pleiotropy. Average causal effect estimates, estimated standard errors, root mean squared errors, empirical Type I error rates (*θ* = 0) and empirical coverage (*θ* = 0.3) for a variety of MR methods.

| Method | Causal Effect $\theta = 0$ | | | | Causal Effect $\theta = 0.3$ | | | |
| --- | --- | --- | --- | --- | --- | --- | --- | --- |
|  | Mean | SE | RMSE | Type I | Mean | SE | RMSE | Coverage |
| IVW | 0.000 | 0.022 | 0.022 | 0.057 | 0.296 | 0.022 | 0.023 | 0.942 |
| MR-Egger | -0.003 | 0.065 | 0.065 | 0.055 | 0.265 | 0.066 | 0.077 | 0.912 |
| Median S | 0.000 | 0.010 | 0.011 | 0.070 | 0.300 | 0.012 | 0.012 | 0.946 |
| Median W | 0.000 | 0.010 | 0.010 | 0.061 | 0.294 | 0.012 | 0.014 | 0.915 |
| Mode S | 0.000 | 0.020 | 0.014 | 0.009 | 0.295 | 0.025 | 0.018 | 0.989 |
| Mode W | 0.000 | 0.016 | 0.012 | 0.011 | 0.288 | 0.020 | 0.019 | 0.954 |
| Lasso | 0.000 | 0.007 | 0.007 | 0.089 | 0.296 | 0.008 | 0.010 | 0.861 |
| MR-Presso | 0.000 | 0.007 | 0.008 | 0.086 | 0.296 | 0.010 | 0.012 | 0.884 |
| MR-Raps | 0.000 | 0.019 | 0.019 | 0.048 | 0.300 | 0.020 | 0.020 | 0.951 |
| ConMix | 0.000 | — | 0.008 | 0.218 | 0.298 | — | 0.011 | 0.778 |
| JAM-MR | 0.000 | 0.007 | 0.008 | 0.074 | 0.294 | 0.009 | 0.013 | 0.851 |
| Oracle | 0.000 | 0.007 | 0.007 | 0.045 | 0.296 | 0.008 | 0.009 | 0.919 |

**Table 3:** Scenario 2: Directional pleiotropy, InSIDE assumption satisfied. Average causal effect estimates, estimated standard errors, root mean squared errors, empirical Type I error rates (*θ* = 0) and empirical coverage (*θ* = 0.3) for a variety of MR methods.

| Method | Causal Effect $\theta = 0$ | | | | Causal Effect $\theta = 0.3$ | | | |
| --- | --- | --- | --- | --- | --- | --- | --- | --- |
|  | Mean | SE | RMSE | Type I | Mean | SE | RMSE | Coverage |
| IVW | 0.208 | 0.019 | 0.208 | 1.000 | 0.503 | 0.019 | 0.204 | 0.000 |
| MR-Egger | 0.000 | 0.056 | 0.058 | 0.067 | 0.264 | 0.057 | 0.068 | 0.910 |
| Median S | 0.068 | 0.012 | 0.069 | 1.000 | 0.384 | 0.014 | 0.085 | 0.000 |
| Median W | 0.052 | 0.011 | 0.053 | 0.999 | 0.358 | 0.013 | 0.059 | 0.008 |
| Mode S | 0.000 | 0.017 | 0.012 | 0.009 | 0.297 | 0.023 | 0.016 | 0.995 |
| Mode W | 0.000 | 0.014 | 0.011 | 0.012 | 0.290 | 0.018 | 0.017 | 0.961 |
| Lasso | 0.028 | 0.007 | 0.029 | 0.948 | 0.339 | 0.008 | 0.041 | 0.021 |
| MR-Presso | 0.077 | 0.011 | 0.078 | 1.000 | 0.396 | 0.013 | 0.098 | 0.000 |
| MR-Raps | 0.170 | 0.019 | 0.171 | 1.000 | 0.472 | 0.020 | 0.173 | 0.000 |
| ConMix | 0.001 | — | 0.008 | 0.183 | 0.302 | — | 0.011 | 0.771 |
| JAM-MR | 0.005 | 0.007 | 0.009 | 0.106 | 0.301 | 0.009 | 0.012 | 0.897 |
| Oracle | 0.000 | 0.007 | 0.006 | 0.044 | 0.296 | 0.008 | 0.009 | 0.919 |

**Table 4:** Scenario 3: Directional pleiotropy, InSIDE assumption violated. Average causal effect estimates, estimated standard errors, root mean squared errors, empirical Type I error rates (*θ* = 0) and empirical coverage (*θ* = 0.3) for a variety of MR methods.

| Method | Causal Effect $\theta = 0$ | | | | Causal Effect $\theta = 0.3$ | | | |
| --- | --- | --- | --- | --- | --- | --- | --- | --- |
|  | Mean | SE | RMSE | Type I | Mean | SE | RMSE | Coverage |
| IVW | 0.372 | 0.022 | 0.372 | 1.000 | 0.668 | 0.023 | 0.369 | 0.000 |
| MR-Egger | 0.705 | 0.060 | 0.707 | 1.000 | 0.978 | 0.062 | 0.680 | 0.000 |
| Median S | 0.069 | 0.012 | 0.070 | 1.000 | 0.387 | 0.015 | 0.088 | 0.000 |
| Median W | 0.167 | 0.018 | 0.175 | 1.000 | 0.496 | 0.021 | 0.204 | 0.000 |
| Mode S | 0.000 | 0.020 | 0.013 | 0.007 | 0.295 | 0.020 | 0.016 | 0.980 |
| Mode W | 0.000 | 0.017 | 0.011 | 0.016 | 0.288 | 0.016 | 0.018 | 0.929 |
| Lasso | 0.045 | 0.008 | 0.047 | 0.996 | 0.366 | 0.009 | 0.068 | 0.000 |
| MR-Presso | 0.204 | 0.017 | 0.206 | 1.000 | 0.517 | 0.018 | 0.218 | 0.000 |
| MR-Raps | 0.338 | 0.024 | 0.339 | 1.000 | 0.640 | 0.024 | 0.340 | 0.000 |
| ConMix | 0.001 | — | 0.009 | 0.220 | 0.299 | — | 0.010 | 0.813 |
| JAM-MR | 0.002 | 0.007 | 0.008 | 0.074 | 0.296 | 0.009 | 0.011 | 0.899 |
| Oracle | 0.000 | 0.007 | 0.007 | 0.053 | 0.296 | 0.008 | 0.009 | 0.919 |

In the first simulation, all Mendelian randomization methods provided nearly unbiased estimates of the causal effect of interest. This was the case even for the standard inverse-variance weighted method, which does not perform pleiotropy adjustments (the IVW was implemented with a random-effects adjustment, hence the fairly accurate standard error estimates). In this “balanced pleiotropy” scenario, the pleiotropic effects cancel out and it is relatively straightforward to obtain an accurate causal effect estimate.

The JAM-MR implementation attained standard errors close to those obtained by the oracle IVW and had one of the smallest mean squared errors among the competing approaches. In terms of Type I error rates for *θ* = 0, most of the competing methods attained nominal levels, with the contamination mixture being the only approach that exhibited significant inflation. For *θ* = 0.3, there was a slight inflation in the coverage of confidence intervals for the Lasso, MR-Presso, contamination mixture and JAM-MR algorithms. Other than that, the performance of the various Mendelian randomization algorithms was similar for *θ* = 0 and for *θ* = 0.3.

In the directional pleiotropy scenario 2 (Table 3), there were larger deviations in the performance of the various methods. Bias in causal effect estimates was observed for the majority of methods, with mode-based estimation, the contamination mixture and JAM-MR being the most accurate algorithms. Among these three approaches, JAM-MR and the contamination mixture attained smaller mean squared errors. The standard errors computed using JAM-MR were smaller than those of the mode and close to those of the oracle estimator. On the other hand, the mode-based approach offered better calibration of Type I error rates and coverage of confidence intervals; in fact, the method is conservative and attains Type I error rates below the 5% threshold. Both JAM-MR and the contamination mixture exhibited a slight inflation in Type I error rates, although the issue was less pronounced for JAM-MR.

Among the other methods, MR-Egger returned fairly accurate causal effect estimates and well-calibrated confidence intervals, but had a significantly higher standard error compared to all the other approaches. The lack of power associated with MR-Egger is well-known in the Mendelian randomization literature (e.g. Slob and Burgess, 2019). The remaining methods were subject to different degrees of bias in causal effect estimates with the Lasso approach being the least biased in this scenario. This inaccuracy also gave rise to large Type I error rates.

A similar pattern of results was observed in scenario 3 (Table 4). JAM-MR, the contamination mixture and mode-based estimation were again the three methods that provided the most accurate causal effect estimates. Among these methods, JAM-MR yielded smaller mean squared error and standard errors close to those from the oracle IVW approach. The contamination mixture offered similar mean squared error to JAM-MR but worse inflation of Type I error rates, while the mode-based approach provided confidence intervals with nominal coverage. Again, the rest of the competing methods were subject to large biases. This included the MR-Egger method, since the InSIDE assumption was violated in this simulation.

The Type I error rates observed for some of the competing methods are rather extreme. These error rates were mainly due to biases in causal effect estimates. Our simulation was rather challenging, as it contained a fairly large number of pleiotropic variants with similar univariate causal effect estimates and this was difficult for some of the methods to tackle. For example, the MR-Presso method relies on outlier removal and can be expected to struggle in simulations where many genetic variants exhibit similar pleiotropic effects. In addition, the fairly large sample sizes used in our simulation study gave rise to quite narrow confidence intervals, and the bias exhibited by these methods implied that the confidence intervals were centered at the wrong estimates, therefore Type I error rates were inflated. Simulations with smaller sample sizes are reported in the Appendix, and Type I error rate inflation for the various methods is slightly smaller in these simulations. Although it is perhaps counterintuitive that some methods may perform better with smaller sample sizes, similar observations have been made elsewhere (e.g. in some of the simulations in Qi and Chatterjee, 2019).

In aaddition, in the Appendix we consider simulations with different numbers of genetic variants, a varying proportion of genetic variation in the risk factor that is explained by the genetic variants and different proportions of pleiotropic variants. In the majority of simulations, JAM-MR was able to consistently estimate the causal effect of interest. Of note was a small deterioration in the algorithm’s performance when both the GWAS sample sizes and the proportion of genetic variation in the risk factor were small. The mode and the contamination mixture remained the best competitors throughout these simulations.

We should also mention that many of the existing methods would perform quite well in different simulation scenarios, and some of them (such as the median and MR-Raps) come with theoretical guarantees of good asymptotic performance. Their poor behaviour here should be taken as an indication that JAM-MR performs well even in challenging simulation scenarios where other methods can struggle, rather than a general-purpose criticism towards these methods.

In the left-hand panel of Figure 3 we visualize the causal effect estimates obtained from the various methods for the directional pleiotropy simulation with the InSIDE assumption satisfied (scenario 2) and *θ* = 0.3. In addition, in the right-hand panel of Figure 3, we have plotted causal effect estimates and confidence intervals obtained by running JAM-MR with various values of *w*. The plot illustrates how the algorithm can be used as a tool for sensitivity analysis. Causal effect estimates for small values of *w* were much larger than those for moderate and large values, indicating that pleiotropic variants are present in the dataset and cause upwards bias if not accounted for. As we increase *w* towards the minimum-standard-error value, plotted in red, causal effect estimates become unbiased. However, for large values of *w* the estimates become more variable and eventually unstable. This is because the pleiotropic loss function removes most of the valid SNPs and causal effect estimation is based on only a small number of genetic variants, increasing standard errors and reducing the power to detect a causal association.

**Figure 3:**
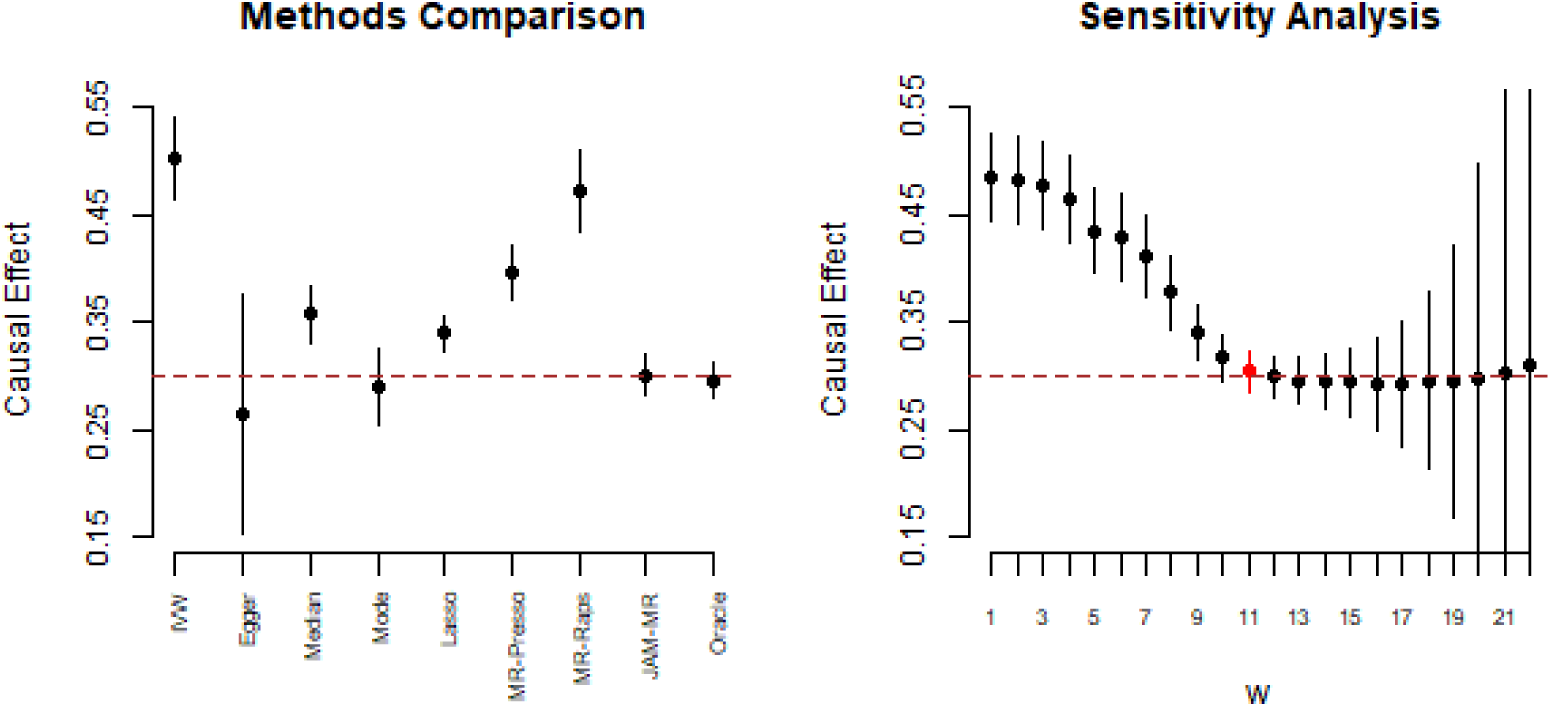
Simulation scenario 2: directional pleiotropy, InSIDE satisfied (*θ* = 0.3). Causal effect estimates and 95% confidence intervals for various Mendelian randomization algorithms (left) and JAM-MR with a range of *w* values (right). The dashed line at 0.3 represents the true causal effect.

The JAM-MR algorithm returns posterior inclusion probabilities for each genetic variant, which can be plotted in a Manhattan plot. In Figure 4 we have plotted inclusion probabilities for a single implementation of the directional pleiotropy simulation scenario 2 with *θ* = 0.3. We used three JAM-MR runs with a small (*w* ≈ 0.03*N*_1_), a moderate (*w* ≈ 0.4*N*_1_) and a large (*w* ≈ 15*N*_1_) value for the tuning parameter. Pleiotropic variants are coloured red. The plot illustrates that our heterogeneity penalization procedure makes accurate selection of the valid instruments, when properly tuned (center panel). When a small value of *w* is used (left panel), the algorithm may retain some pleiotropic SNPs in the analysis. On the other hand, when *w* is large (right panel), the algorithm will correctly downweight pleiotropic variants but will also downweight some of the valid SNPs.

**Figure 4:**
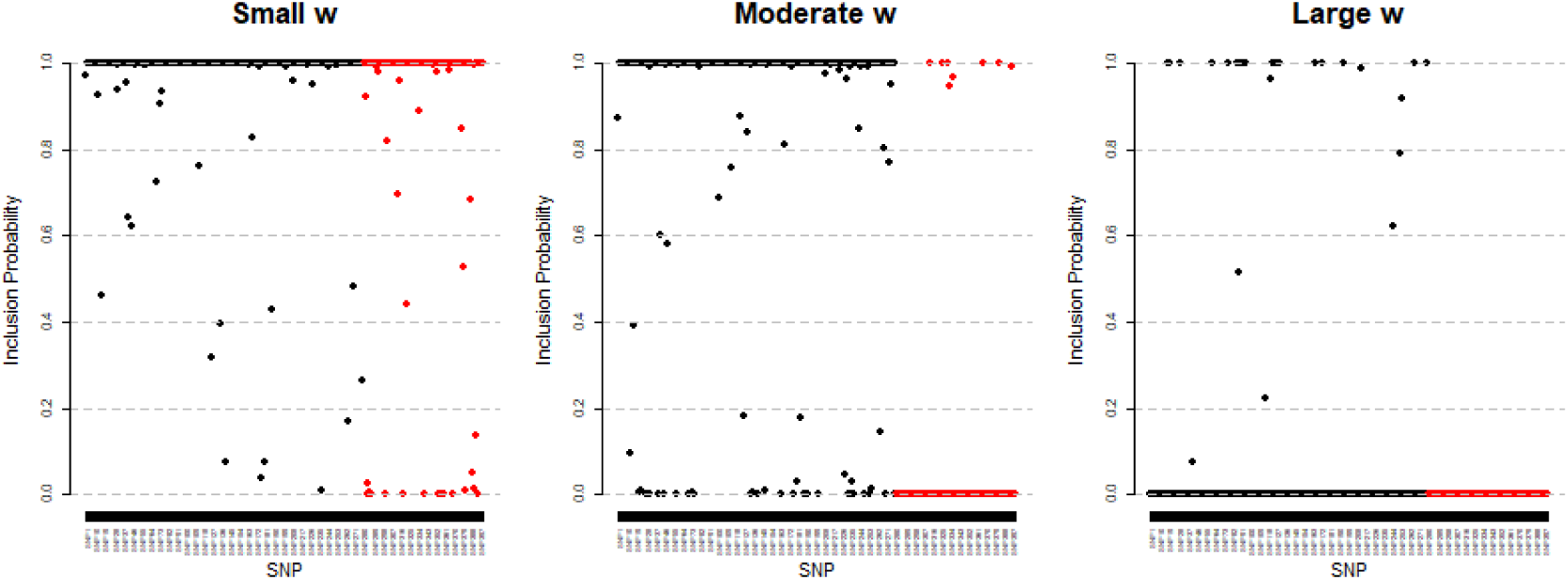
Manhattan plots illustrating the posterior inclusion probabilities assigned to each SNP by the JAM-MR algorithm with small (left), moderate (center) and large (right) values of *w*, for a single implementation of simulation scenario 2 with *θ* = 0.3.

In our simulations, we used a 51-point grid search in order to tune the JAM-MR algorithm. In practice, it may be unnecessary to use such a large grid -often, even a grid as small as 5-10 values will provide a reasonable fit.To verify that, we compared the causal effect estimates obtained from JAM-MR using a 51-point grid search to those obtained using a smaller grid with only 9 points, nested within the 51-point grid. The resulting causal effect estimates were quite similar for all simulation scenarios. In particular, the mean absolute difference in causal effect estimates obtained by the two grid-search procedures across all our simulations was 0.0029, and the mean absolute difference in standard error estimates was 0.00047.

The time required to implement the JAM-MR algorithm is longer than that for other Mendelian randomization approaches. For the simulated datasets with *P* = 400 genetic variants, it takes about 3-5 minutes to implement the JAM-MR algorithm for a single value of *w*. A grid search over many *w* values takes longer, but can be implemented more efficiently if researchers have access to cluster computing facilities, as runs for different *w* values can be executed in parallel.

Overall, the JAM-MR algorithm was one of the best-performing methods in our simulations. The algorithm proved to be robust to a variety of pleiotropic patterns. It yielded practically unbiased causal effect estimates, standard errors similar to those obtained from an oracle IVW estimator and fairly small mean squared errors. Together with mode-based estimation it exhibited the smallest bias in our simulations, and it also attained the smallest mean squared error. A slight overestimation of Type I error rates was observed in some of the simulations, but the issue is less pronounced than for many existing Mendelian randomization algorithms. The most promising competing approaches were mode-based estimation and the contamination mixture. JAM-MR exhibited smaller mean squared errors than the mode, which produced better-calibrated confidence intervals in exchange. The contamination mixture yielded similar mean squared errors as JAM-MR but with a slightly larger Type I error rate inflation.

## 4 Application: causal effect of blood pressure on CHD risk

### 4.1 Blood pressure traits and associated genetic variants

We now illustrate the use of the JAM-MR algorithm in a real-data application. We conduct a Mendelian randomization analysis to assess the effect of blood pressure on the risk of coronary heart disease (CHD). This application has been studied in the past (Bowden et al., 2015; Warren et al., 2017; Bowden et al., 2018) and it is generally accepted that high blood pressure has an increasing effect on the risk of suffering from coronary heart disease, despite the fact that a previous Mendelian randomization analysis (Bowden et al., 2015) did not identify a causal relationship.

Our analysis is novel not in the question it aims to answer but in the data sources it uses. We utilized a recently published meta-GWAS study (Evangelou et al., 2018) which identified hundreds of genetic variants associated with blood pressure. The authors meta-analyzed data from the International Consortium for Blood Pressure (ICBP), the UK BioBank, the US Million Veterans Project and the Estonian Genome Center Biobank (EGCUT). In total, a sample of approximately 1 million individuals of European descent was analyzed. The study confirmed previously reported findings about 258 known and 92 reported but not validated genetic variants associated with blood pressure. It also identified a total of 535 novel associations.

We used two blood pressure traits for our analysis, namely systolic and diastolic blood pressure. For each trait, we used all 258 genetic variants with an already established relation to blood pressure, as well as any of the reported-but-not-validated and novel variants reported to be associated with that trait. Among the novel findings, we excluded from consideration variants which were associated with blood pressure in the “discovery” GWAS but not in the “replication” GWAS in Evangelou et al. (2018). This resulted in a total of *P*_1_ = 395 genetic variants for systolic and *P*_2_ = 391 genetic variants for diastolic blood pressure; our analysis was therefore based on larger numbers of genetic variants than previous Mendelian randomization investigations.

Genetic associations with systolic and diastolic blood pressure were obtained from the Supplementary Tables of Evangelou et al. (2018). We used estimates based on the ICBP dataset of *N*_1_ = 299024 individuals, as this was the only dataset for which genetic associations were reported for all variants. Since the genetic associations with blood pressure were replicated in independent datasets in Evangelou et al. (2018), winner’s curse bias is unlikely to have had a serious effect in our analysis. For the selected variants, we obtained genetic associations with coronary heart disease risk from the CARDIoGRAM-plusC4D Consortium(CARDIoGRAMplusC4D Consortium, 2015), based on a sample of *N*_2_ = 184305 individuals. The variants were mostly independent, as a result of LD pruning in Evangelou et al. (2018).

### 4.2 Mendelian randomization analysis

We implemented JAM-MR and the competing Mendelian randomization methods to identify the causal effect of systolic and diastolic blood pressure on CHD risk. The results are listed in Table 5, where we report the estimated causal effect (log-odds ratio of increase in CHD risk per mmHg increase in blood pressure measurement) and its 95% confidence interval for each of the two traits and each Mendelian randomization method. Confidence intervals for the JAM-MR algorithm were computed using the truncated random-effects approach. A graphical illustration of the results is provided in Figure 5. Tables of summary statistics and JAM-MR inclusion probabilities for each SNP can be found in the supplementary material, and the corresponding Manhattan plots are provided in the Appendix.

**Table 5:** Log-odds ratios of increase in CHD risk per 1mmHg increase in the corresponding blood pressure measurement. Causal effect estimates and 95 % confidence intervals for a variety of MR methods.

| Method | Systolic Blood Pressure |  | Diastolic Blood Pressure |  |
| --- | --- | --- | --- | --- |
|  | Estimate | 95% C.I. | Estimate | 95% C.I. |
| IVW | 0.034 | (0.026 , 0.041) | 0.048 | (0.035 , 0.061) |
| MR-Egger | 0.024 | (0.008 , 0.041) | 0.083 | (0.058 , 0.109) |
| Median S | 0.038 | (0.030 , 0.046) | 0.053 | (0.038 , 0.068) |
| Median W | 0.032 | (0.024 , 0.040) | 0.060 | (0.047 , 0.073) |
| Mode S | 0.026 | (-0.087 , 0.139) | 0.061 | (-0.059 , 0.181) |
| Mode W | 0.026 | (-0.087 , 0.139) | 0.061 | (-0.057 , 0.179) |
| Lasso | 0.037 | (0.032 , 0.042) | 0.048 | (0.040 , 0.057) |
| MR-Presso | 0.038 | (0.032 , 0.045) | 0.052 | (0.042 , 0.063) |
| MR-Raps | 0.039 | (0.032 , 0.045) | 0.055 | (0.043 , 0.066) |
| ConMix | 0.038 | (0.031 , 0.043) | 0.051 | (0.040 , 0.063) |
| JAM-MR | 0.039 | (0.030 , 0.047) | 0.067 | (0.054 , 0.079) |

**Figure 5:**
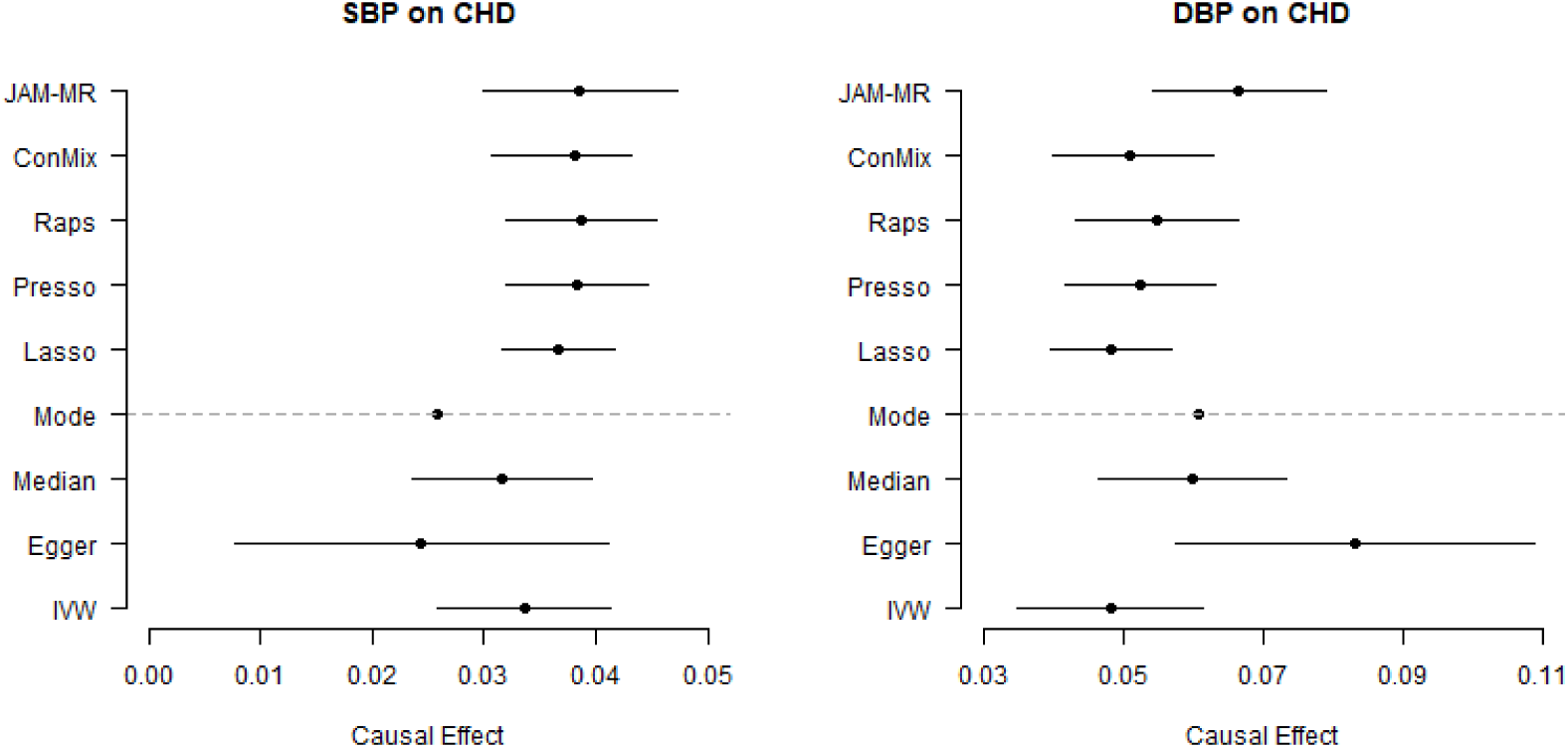
Log-odds ratios of increase in CHD risk per 1mmHg increase in systolic and diastolic blood pressure. Point estimates and 95% confidence intervals for various Mendelian randomization methods.

The reported effects from the various methods confirm that increased blood pressure, both systolic and diastolic, has a deleterious effect on coronary heart disease risk. For systolic blood pressure, the causal effect reported by the various methods was in the region of 0.024 − 0.040, corresponding to an odds ratio of *e*^0.024^ = 1.024 to *e*^0.04^ = 1.041. All methods were able to reject the null causal hypothesis at a 95% significance threshold, except the mode-based estimator whose standard error was unusually large. JAM-MR estimates were very close to those reported by other methods. The minimum causal standard error criterion suggested using an implementation with *w* ≈ 0.5*N*_1_, for which the corresponding JAM-MR log-odds ratio estimate was 0.039 (95% confidence interval: (0.030, 0.047)).

For diastolic blood pressure, most of the established Mendelian randomization methods reported causal effect estimates between 0.048 and 0.061, corresponding to odds ratios between 1.049 and 1.063. The JAM-MR estimates were slightly larger, yielding a a log-odds ratio of 0.067 (95% confidence interval (0.054, 0.079)) and an effect estimate of *e*^0.067^ = 1.069. These were attained for *w* ≈ 0.29*N*_1_. This difference is mainly due to JAM-MR downweighting genetic variants that are weak instruments; the estimate of 0.067 is close to what would be obtained by running the competing methods using only the 258 genetic variants with established associations with blood pressure. Once again, all methods were able to reject the null causal hypothesis at a 95% level, except the mode-based estimator.

### 4.3 Interpretation of results

The two JAM-MR implementations assigned posterior inclusion probabilities larger than 0.5 to 142 genetic variants for systolic and 169 variants for diastolic blood pressure. Of the 253 variants for systolic blood pressure that were assigned posterior probability lower than 0.5, 116 had univariate causal effect estimates below the overall reported value of 0.037 and 137 had univariate estimates above that value. For diastolic blood pressure, there were 222 variants assigned posterior probabilities lower than 0.5, 121 of which had smaller and 101 had larger univariate ratio estimates than the reported value of 0.067. Not all genetic variants were downweighted due to pleiotropy; some were assigned low posterior probabilities because their association with the two blood pressure traits were not strong enough. Nevertheless, the downweighting of genetic variants both larger and smaller than the overall causal effect estimates in almost equal numbers suggests a fairly balanced pleiotropic pattern.

To further explore the effect of pleiotropic variants on the estimated effect of blood pressure on coronary heart disease risk, we can use the sensitivity analysis plots of Figure 6. The overall impact of pleiotropy in this application seems rather small, as similar causal effect estimates are obtained for a wide range of *w* values. There is perhaps a slight downward bias for diastolic blood pressure with small *w* values, while JAM-MR estimates with unnecessarily large *w* become unstable, as in the simulations.

**Figure 6:**
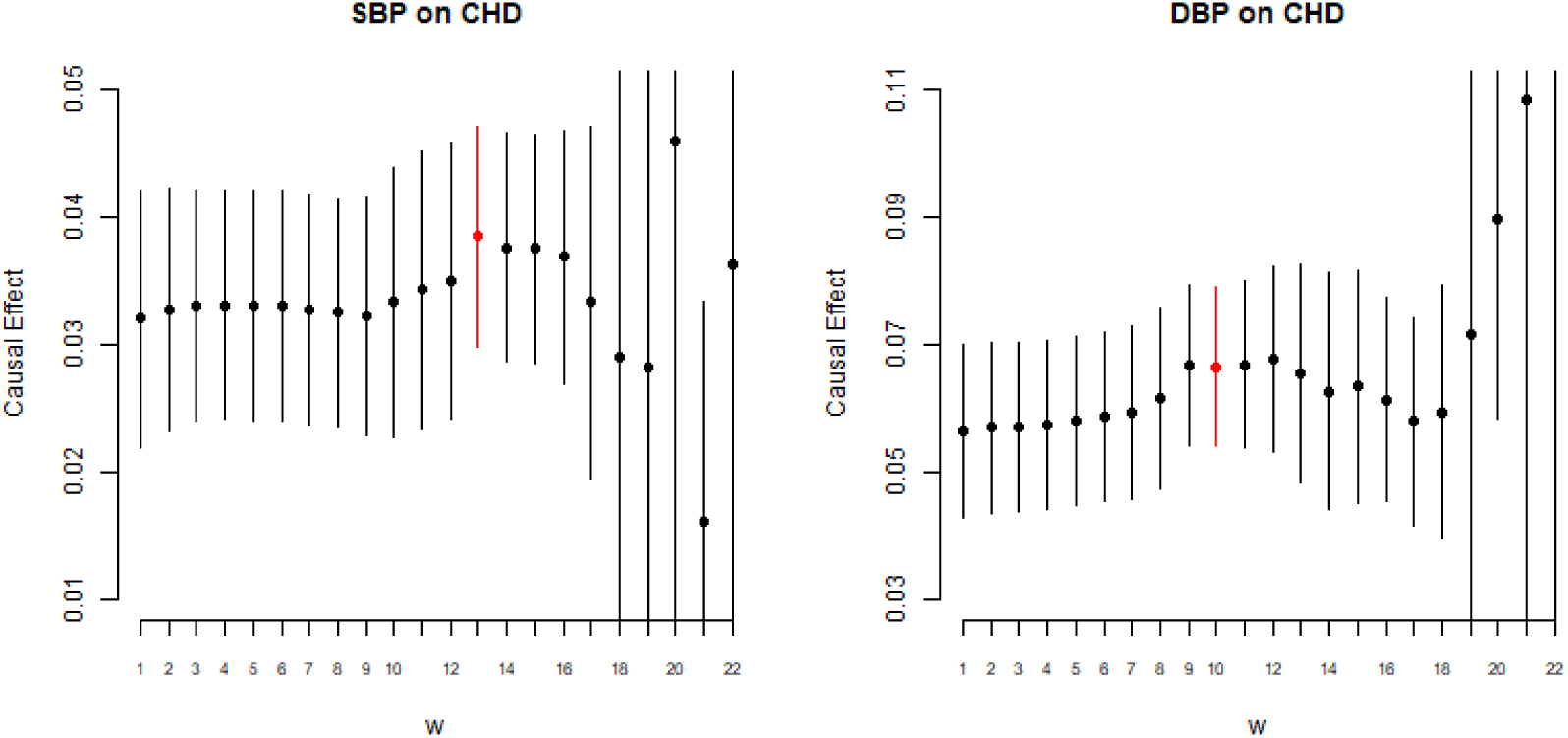
Log-odds ratios of increase in CHD risk per 1mmHg increase in systolic and diastolic blood pressure. Point estimates and 95% confidence intervals for JAM-MR implementations with various values of the tuning parameter.

The MR-Egger method rejects the null hypothesis of no causal association and indicates the existence of a causal effect on CHD risk for both blood pressure traits. MR-Egger was the method used to generate the null findings in Bowden et al. (2015). This seeming inconsistency is due to the larger sample sizes and numbers of SNPs that we have used in this paper. The analysis in Bowden et al. (2015) was based only on 29 genetic variants that were known to be associated with blood pressure at that point, and its power to detect a causal association was lower.

The mode-based estimation method was implemented using the default function in the R package `MendelianRandomization`. The method yielded much larger standard errors than other algorithms. This was due to the presence of a few genetic variants with extreme outlying effects in our dataset, which were in turn the result of weak instrument bias. The presence of of weak instruments can have adverse effects on the performance of the mode-based approach. In fact, the effects can be more severe than those reported in Table 5. The estimates for diastolic blood pressure in Table 5 were obtained after discarding three obvious outliers from the dataset. Using all 391 genetic variants, the resulting causal effect estimate was −0.063 (95% confidence interval (−22.27, 22.152)) for the simple and −0.063 (95% confidence interval (−22.260, 22.134)) for the weighted version of the mode-based method; both confidence intervals were too wide to be of practical use. This poor performance of the method was not observed in our simulation study because our simulation design did not generate very weak instruments.

## 5 Discussion

In this paper, we have developed a new algorithm for causal effect estimation in Mendelian randomization when some of the candidate instruments are pleiotropic. Our algorithm uses Bayesian variable selection to identify sets of genetic variants with homogeneous causal effect estimates, and model averaging to account for uncertainty in the selection of instruments to use. A wide range of simulation studies and a real-data application using data from a recent GWAS to study the effect of blood pressure on coronary heart disease risk demonstrate how using model averaging to account for uncertainty in variable selection leads to robsut causal inference.

Compared to other approaches for Mendelian randomization with pleiotropic variants, the JAM-MR algorithm has a number of attractive features. The use of model averaging allows many variants to have small contributions to the overall causal effect estimate and offers uncertainty quantification for genetic associations with the risk factor. Unlike the competing approaches discussed in this paper, our algorithm uses the Bayesian framework. Although not explored in this paper, this raises the possibility of incorporating prior information on the biological function of SNPs or genes, something we intend to explore in the future. JAM-MR’s stochastic search procedure is quite flexible and can explore large parameter spaces of causal configurations; as a result, the algorithm is quite efficient when used with large numbers of genetic variants. Our algorithm also provides a natural framework for incorporating genetic correlations into a Mendelian randomization analysis and selecting the most relevant variants from a densely genotyped region. Common approaches for Mendelian randomization typically assume that genetic variants are independent. In related work, we are investigating the advantages of utilizing JAM’s variable selection compared to pruning and other approaches for Mendelian randomization with correlated instruments, in order to incorporate multiple correlated effects from regions which harbour complex genetic signals for the trait of interest.

In practice the JAM-MR algorithm performs particularly well when implemented in large-scale GWAS studies with hundreds of genetic variants that can explain a large proportion of variation in the risk factor, as demonstrated in our simulation study.

Limitations of our approach include the algorithm’s imperfect calibration of standard errors. Thankfully, the inflation of Type I error rates is not severe. A heuristic yet practical adjustment could be that researchers report 99% confidence intervals instead of the traditional 95% threshold, when using the JAM-MR algorithm. It would also be useful to devise an efficient automatic procedure for specifying the tuning parameter *w* that does not have to rely on a grid search. Running JAM-MR with several *w* values can help visualize how pleiotropic variants affect the causal effect estimate, but comes with an increased computational cost and it would be desirable to obtain an accurate causal effect estimate based on a single implementation of the algorithm. JAM-MR could also benefit from a more elaborate implementation of the reversible-jump procedure, for example using parallel tempering. Finally, another extension would be to construct a fully Bayesian version of the algorithm.

In conclusion, JAM-MR performs pleiotropy-robust causal effect estimation for Mendelian randomization. Our algorithm has a number of desirable features, exhibits good performance in simulations and has been used to implement a Mendelian randomization analysis of the effect of blood pressure on CHD risk, using a recent large-scale blood pressure meta-GWAS. We therefore hope that JAM-MR will become a valuable addition to the Mendelian randomization literature.

The JAM-MR algorithm has been implemented in `R` as part of the GitHub package `R2BGLiMS`, available at https://github.com/pjnewcombe/R2BGLiMS.

## 6 Acknowledgements

This work was supported by the UK Medical Research Council (Core Medical Research Council Biostatistics Unit Funding Code: MC UU 00002/7). Apostolos Gkatzionis and Paul Newcombe were supported by a Medical Research Council Methodology Research Panel grant (Grant Number RG88311). Stephen Burgess was supported by a Sir Henry Dale Fellowship jointly funded by the Wellcome Trust and the Royal Society (Grant Number 204623/Z/16/Z). David Conti was supported by a grant from the National Cancer Institute (Grant Number NIH/NCI P01CA196569).

## Appendix

### Simulation design

Figure 7 illustrates the design of our simulation study. We plot a histogram of the negative logarithms of univariate p-values generated for each genetic variant (left) and a histogram of univariate causal effect estimates obtained from all the simulated genetic variants (right). The two histograms are constructed using data from all 1000 replications of simulation scenario 2 with *θ* = 0. Similar plots were obtained for the other two simulation scenarios, as well as for *θ* = 0.3.

**Figure 7:**
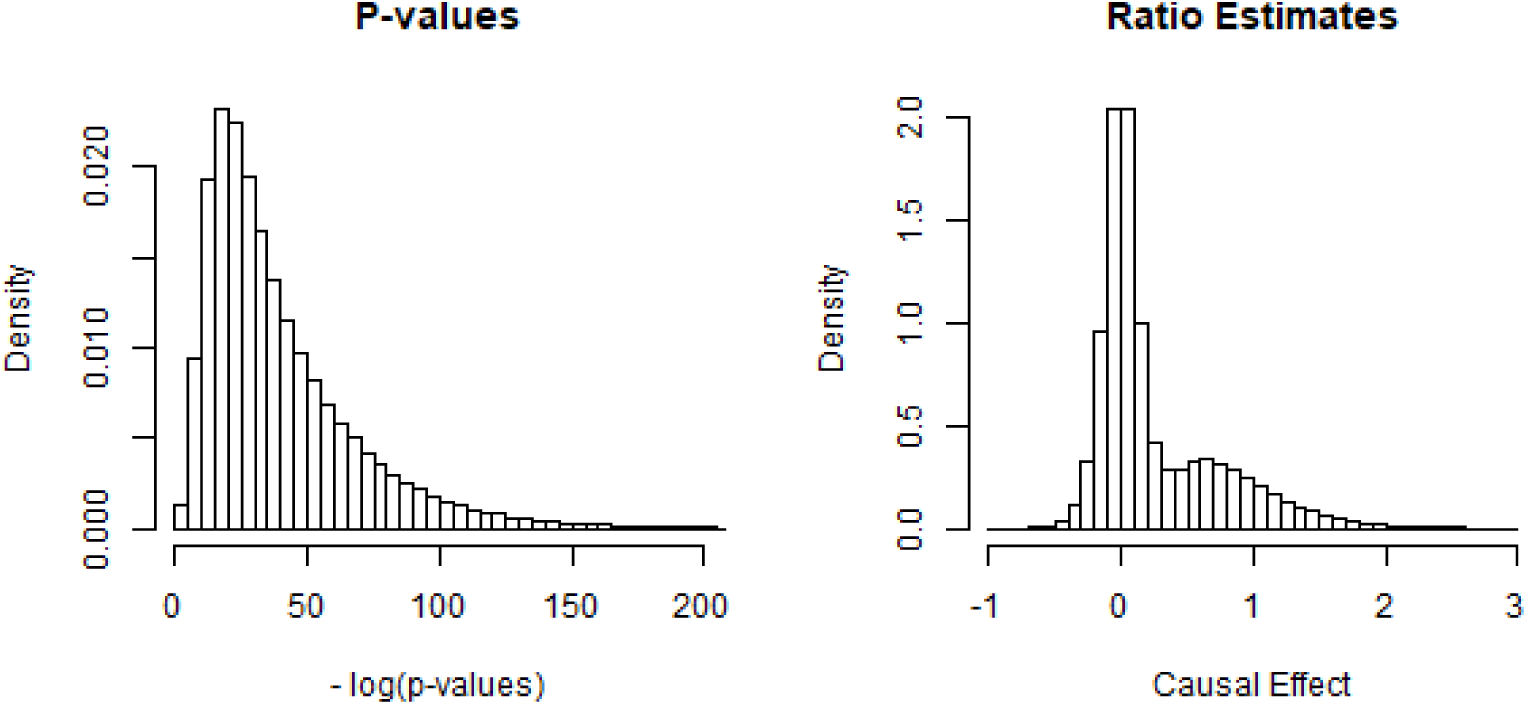
Histogram of univariate p-values of association between each genetic variant and the risk factor (left) and univariate causal effect estimates (right) over all replications of simulation scenario 2 with *θ* = 0.

**Figure 8:**
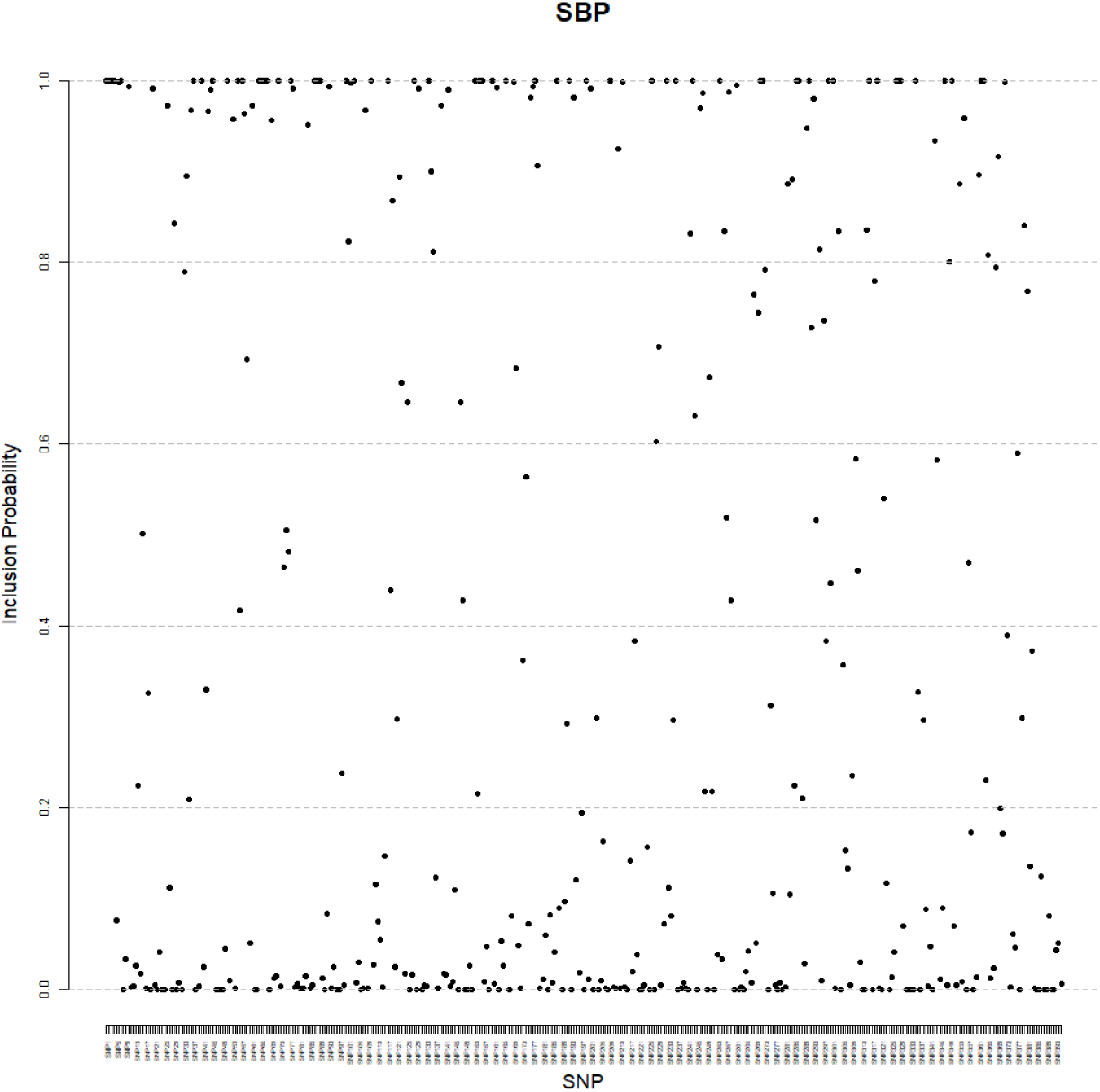
Manhattan plot of posterior inclusion probabilities for the 395 SBP-associated genetic variants.

**Figure 9:**
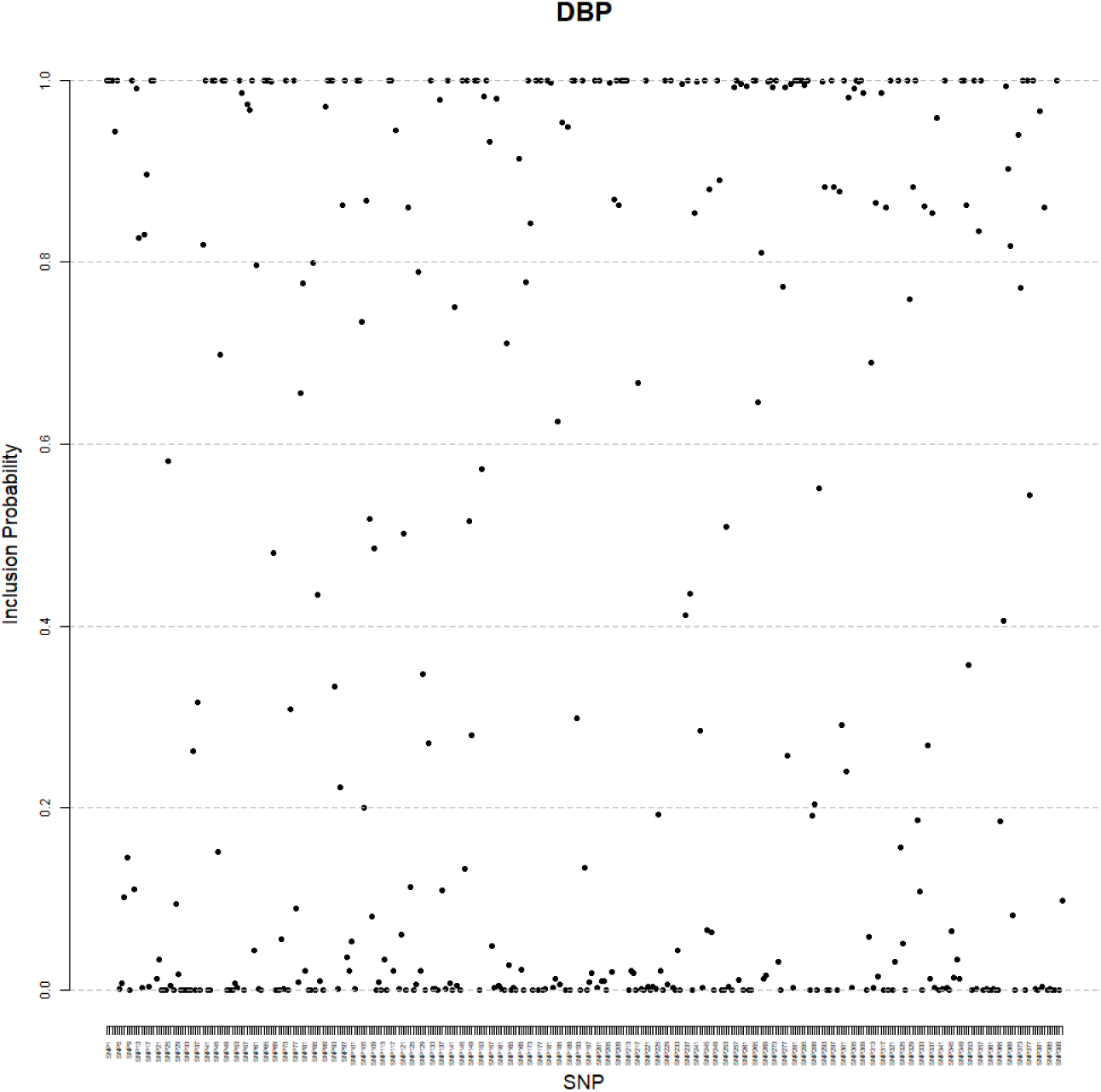
Manhattan plot of posterior inclusion probabilities for the 391 DBP-associated genetic variants.

### Competing methods

Mendelian randomization in the presence of invalid instruments is a very active area of research. A wide range of statistical techniques have been used to either identify the pleiotropic variants and remove them from the analysis, or robustify the process of causal effect estimation. Here we give a brief description of the methods used in our simulations.

One of the earliest and most widely used approaches is MR-Egger regression (Bowden et al., 2015). Like the inverse-variance weighted estimate, MR-Egger regression is motivated from the meta-analysis literature. Loosely, it consists of fitting a linear regression of the SNP-outcome association estimates on the SNP-exposure estimates: 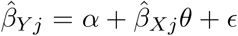.

The IVW estimate is obtained when *α* = 0. When *α* ≠ 0, the intercept models the aggregate pleiotropic contribution of all SNPs on the outcome. MR-Egger yields consistent causal effect estimates under the assumption that instrument strength is independent of the genetic variants’ direct effects on the outcome (InSIDE). However, as illustrated in our simulation study, the method’s power to detect causal associations is often quite low.

Another common approach is to estimate the causal effect of interest by the (weighted or unweighted) median of univariate estimates for all available genetic variants. This median estimator is robust to outlying pleiotropic effects and is asymptotically unbiased if more than 50% of the available genetic variants are valid instruments.

Mode-based estimation (Hartwig et al., 2016) fits a kernel density to the univariate causal effect estimates and uses the mode of that density as the overall estimate; standard errors are then computed based on a bootstrap procedure. This approach weakens the majority assumption of the median and yields accurate causal effect estimates under the assumption that only a plurality of genetic instruments are valid.

Lasso regularization has also been considered in order to identify which genetic variants exhibit pleiotropic effects. This was proposed by Kang et al. (2016) in the presence of individual-level data and adapted by Rees et al. (2019) to situations where only summary-level data are available. Rees et al. (2019) also developed robust regression techniques for causal effect estimation.

Another approach that relies on variable selection is MR-Presso Verbanck et al. (2018), which performs outlier detection and deletion by conducting a hypothesis test of pleiotropy for each genetic variant.

The contamination mixture approach Burgess et al. (2019) uses a mixture model for the univariate causal effect estimates. The mixture contains two components: valid instruments are assumed to be normally distributed around the true causal effect and invalid instruments are normally distributed around zero. The method constructs a likelihood based on that mixture model, and the likelihood is maximized over both parameter values and assignments of SNPs to the two mixture components; the maximization relies on profile likelihood techniques.

The MR-Raps method Zhao et al. (2018) assumes a random-effects distribution for pleiotropic effects, and also relies on profile likelihood and robust regression techniques for causal effect estimation.

For an extensive comparison of pleiotropy-robust Mendelian randomization algorithms using summary data, we refer the reader to two recent review papers (Slob and Burgess, 2019; Qi and Chatterjee, 2019).

The list of methods that we considered in our simulation study is not exhaustive. For example, we did not implement the heterogeneity penalization approach presented in Burgess et al. (2018). Similar to JAM-MR, this approach relies on model averaging, but instead of our algorithm’s stochastic search it uses an exhaustive search over possible models, and is therefore only applicable for small numbers of genetic variants. We have also not considered the mixture modelling method of Qi and Chatterjee (2018); this method was shown to perform well in terms of Type I error calibration in a recent study (Qi and Chatterjee, 2019), but in order to function efficiently it also requires access to summary statistics for additional genetic variants throughout the genome that are not associated with the risk factor. We did not generate such variants in our simulations, since it would have dramatically increased their computational cost. Finally, we did not utilize methods requiring the availability of individual-level data, such as sisVIVE (Kang et al., 2016) or the instrumental variables method of Koop et al. (2012) which also uses Reversible-Jump MCMC.

### Implementation of the various methods in the simulation study

For the implementation of the various Mendelian randomization methods in our simulation study, we used available `R` packages. In particular, we used the packages `MendelianRandomization` (for the IVW, MR-Egger, contamination mixture, median and mode-based methods), `MRPRESSO` (for MR-Presso) and `mr.raps` (for MR-Raps). For the Lasso method, we used the R code provided in the Appendix of Rees et al. (2019).

For the median and mode-based methods we implemented both an unweighted and a weighted version. For the lasso method we used the “heterogeneity” approach described in Rees et al. (2019) to specify a value for the tuning parameter. MR-Raps was implemented using the Tukey loss function and an overdispersed model. For the other methods, we used the default settings in the corresponding `R` packages.

For the implementation of JAM-MR, we used a grid search consisting of 50 points, evenly scaled on a logarithmic scale from *w* = 0.01*N*_1_ to *w* = 100*N*_1_, plus an execution for *w* = 0 (the original JAM algorithm with no pleiotropy penalization). In each instance, the algorithm was run for 1 million iterations. Since genetic variants were assumed to be independent, we did not require a reference dataset for the matrix of genetic correlations. Instead, the matrix *G*^*T*^ *G* was set as follows. The off-diagonal elements were set equal to zero, reflecting our assumption of independent genetic variants. The diagonal elements were computed by 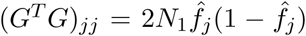, where 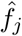 is the estimated effect allele frequency for variant *G*_*j*_. Finally, JAM-MR’s model-specific estimates were computed using the truncated multiplicative random effects approach.

### Comparison between approaches for computing JAM-MR model-specific estimates

Here we compare three approaches to computing model-specific estimates and standard errors when averaging over the models visited by JAM-MR to compute an overall causal effect estimate. The three approaches to be compared are (a) the standard inverse-variance weighted estimate, (b) the IVW estimate using a multiplicative random-effects assumption to compute the standard error, and (c) the truncated normal multiplicative random-effects model described in Section 2.

Each of the three approaches was implemented in the simulation scenarios of Section 3 and causal effect estimates and standard errors were obtained. The results are summarized in Table 6 for all three simulation scenarios. We report average causal effect estimates, estimated standard errors, root mean squared errors and Type I error rates.

**Table 6:** Comparison between the fixed-effects (”IVW”), multiplicative random-effects (”Mult”) and truncated multiplicative random-effects (”Trunc”) approaches for computing model-specific JAM-MR causal effect estimates.

| Method | $\theta = 0$ | | | | $\theta = 0.3$ | | | |
| --- | --- | --- | --- | --- | --- | --- | --- | --- |
|  | Mean | SE | RMSE | Type I | Mean | SE | RMSE | Type I |
| Balanced Pleiotropy |  |  |  |  |  |  |  |  |
| IVW | 0.000 | 0.006 | 0.022 | 0.633 | 0.296 | 0.007 | 0.023 | 0.581 |
| Mult | 0.000 | 0.007 | 0.009 | 0.134 | 0.294 | 0.008 | 0.013 | 0.256 |
| Trunc | 0.000 | 0.007 | 0.008 | 0.074 | 0.294 | 0.009 | 0.013 | 0.149 |
| Directional Pleiotropy, InSIDE Satisfied |  |  |  |  |  |  |  |  |
| IVW | 0.206 | 0.006 | 0.207 | 1.000 | 0.502 | 0.007 | 0.202 | 1.000 |
| Mult | 0.007 | 0.007 | 0.011 | 0.240 | 0.305 | 0.008 | 0.012 | 0.208 |
| Trunc | 0.005 | 0.007 | 0.009 | 0.106 | 0.301 | 0.009 | 0.012 | 0.103 |
| Directional Pleiotropy, InSIDE Violated |  |  |  |  |  |  |  |  |
| IVW | 0.372 | 0.005 | 0.372 | 1.000 | 0.669 | 0.006 | 0.369 | 1.000 |
| Mult | 0.003 | 0.007 | 0.009 | 0.126 | 0.298 | 0.008 | 0.011 | 0.137 |
| Trunc | 0.002 | 0.007 | 0.008 | 0.074 | 0.296 | 0.009 | 0.011 | 0.101 |

The table illustrates the benefit provided by using the truncated approach, as it offers unbiased causal effect estimation and reasonable Type I error rates. The multiplicative random-effects approach is also able to accurately estimate the causal effect but has inflated Type I error rates in comparison. On the other hand, using the default IVW approach yields poor results. This is because the IVW approach does not combine well with JAM-MR’s minimum-standard-error tuning process. When implementing JAM-MR with the standard IVW model-specific estimates, the run with *w* = 0 is often selected as the one with the smallest causal standard error. For *w* = 0, the algorithm implements no pleiotropy penalization and biased estimates are obtained. If we fix *w* instead of implementing a grid search, the fixed-effects IVW method performs similar to the multiplicative random-effects approach -in fact, the two methods give the same causal effect estimate (Burgess and Bowden, 2016). In any case, truncation is the most promising approach.

### Additional simulations

In this section we report simulation results from additional simulations, not included in the main part of the paper. To assess the robustness of JAM-MR and other Mendelian randomization algorithms to a variety of simulation scenarios, we performed two additional sets of simulations. First, we modified the sample sizes of the two GWAS studies, the number of genetic variants included in the Mendelian randomization analysis and the proportion of variation in the risk factor explained by the genetic variants. Specifically, we considered three settings:

- *N*_1_ = *N*_2_ = 50000 individuals in each GWAS and *P* = 50 genetic variants that explain 3% of the genetic variation in the risk factor.
- *N*_1_ = *N*_2_ = 100000 individuals in each GWAS and *P* = 100 genetic variants that explain 5% of the genetic variation in the risk factor.
- *N*_1_ = *N*_2_ = 200000 individuals in each GWAS and *P* = 200 genetic variants that explain 7% of the genetic variation in the risk factor.

In the second set of simulations, we modified the proportion of genetic variants with pleiotropic effects. We used the default simulation settings (*N*_1_ = *N*_2_ = 300000, *P* = 400 and 10% variation in the risk factor explained) and generated two sets of simulations, one with 20% of the genetic variants being invalid and another with 50% of the genetic variants being invalid.

To reduce the computational burden, we only implemented 500 replications of each simulation. The results of these simulations are reported in Tables 7-11. For brevity of presentation, we have only included the weighted versions of the median and the mode.

**Table 7:** Simulation results with sample sizes of *N*_1_ = *N*_2_ = 50000 and *P* = 50 genetic variants that explain 3% genetic variation in the risk factor.

| Method | Causal Effect $\theta = 0$ | | | | Causal Effect $\theta = 0.3$ | | | |
| --- | --- | --- | --- | --- | --- | --- | --- | --- |
|  | Mean | SE | RMSE | Type I | Mean | SE | RMSE | Coverage |
| Balanced Pleiotropy |  |  |  |  |  |  |  |  |
| IVW | 0.005 | 0.064 | 0.064 | 0.054 | 0.287 | 0.066 | 0.071 | 0.916 |
| MR-Egger | -0.010 | 0.181 | 0.185 | 0.068 | 0.220 | 0.188 | 0.214 | 0.920 |
| Median | 0.001 | 0.044 | 0.046 | 0.060 | 0.281 | 0.052 | 0.059 | 0.912 |
| Mode | 0.000 | 0.066 | 0.047 | 0.016 | 0.264 | 0.078 | 0.070 | 0.958 |
| Lasso | 0.002 | 0.029 | 0.044 | 0.192 | 0.288 | 0.033 | 0.058 | 0.742 |
| MR-Presso | 0.002 | 0.036 | 0.046 | 0.124 | 0.288 | 0.044 | 0.061 | 0.830 |
| MR-Raps | 0.005 | 0.061 | 0.060 | 0.060 | 0.297 | 0.065 | 0.066 | 0.936 |
| ConMix | 0.001 | — | 0.042 | 0.078 | 0.299 | — | 0.061 | 0.888 |
| JAM-MR | 0.000 | 0.046 | 0.057 | 0.094 | 0.278 | 0.054 | 0.069 | 0.870 |
| Oracle | 0.001 | 0.031 | 0.029 | 0.040 | 0.289 | 0.038 | 0.037 | 0.958 |
| Directional Pleiotropy, InSIDE Satisfied |  |  |  |  |  |  |  |  |
| IVW | 0.205 | 0.057 | 0.208 | 1.000 | 0.497 | 0.061 | 0.202 | 0.030 |
| MR-Egger | 0.004 | 0.161 | 0.169 | 0.066 | 0.220 | 0.168 | 0.194 | 0.916 |
| Median | 0.085 | 0.046 | 0.099 | 0.430 | 0.387 | 0.055 | 0.104 | 0.674 |
| Mode | 0.011 | 0.066 | 0.044 | 0.016 | 0.293 | 0.075 | 0.058 | 0.968 |
| Lasso | 0.074 | 0.029 | 0.086 | 0.654 | 0.389 | 0.034 | 0.103 | 0.344 |
| MR-Presso | 0.116 | 0.041 | 0.124 | 0.804 | 0.434 | 0.049 | 0.143 | 0.210 |
| MR-Raps | 0.177 | 0.059 | 0.181 | 0.958 | 0.485 | 0.063 | 0.190 | 0.082 |
| ConMix | 0.010 | — | 0.045 | 0.102 | 0.351 | — | 0.131 | 0.774 |
| JAM-MR | 0.064 | 0.069 | 0.120 | 0.270 | 0.368 | 0.062 | 0.110 | 0.684 |
| Oracle | -0.001 | 0.031 | 0.030 | 0.044 | 0.291 | 0.038 | 0.038 | 0.954 |
| Directional Pleiotropy, InSIDE Violated |  |  |  |  |  |  |  |  |
| IVW | 0.365 | 0.065 | 0.368 | 1.000 | 0.656 | 0.068 | 0.360 | 0.000 |
| MR-Egger | 0.629 | 0.175 | 0.654 | 0.924 | 0.876 | 0.185 | 0.605 | 0.150 |
| Median | 0.262 | 0.058 | 0.295 | 0.932 | 0.570 | 0.066 | 0.299 | 0.114 |
| Mode | 0.012 | 0.066 | 0.058 | 0.018 | 0.297 | 0.096 | 0.081 | 0.968 |
| Lasso | 0.115 | 0.032 | 0.128 | 0.844 | 0.444 | 0.037 | 0.164 | 0.164 |
| MR-Presso | 0.229 | 0.051 | 0.239 | 0.988 | 0.546 | 0.055 | 0.255 | 0.016 |
| MR-Raps | 0.341 | 0.070 | 0.345 | 1.000 | 0.641 | 0.073 | 0.345 | 0.000 |
| ConMix | 0.012 | — | 0.097 | 0.072 | 0.409 | — | 0.291 | 0.822 |
| JAM-MR | 0.261 | 0.062 | 0.380 | 0.536 | 0.575 | 0.067 | 0.383 | 0.358 |
| Oracle | 0.001 | 0.031 | 0.029 | 0.024 | 0.291 | 0.038 | 0.036 | 0.952 |

**Table 8:** Simulation results with sample sizes of *N*_1_ = *N*_2_ = 100000 and *P* = 100 genetic variants that explain 5% genetic variation in the risk factor.

| Method | Causal Effect $\theta = 0$ | | | | Causal Effect $\theta = 0.3$ | | | |
| --- | --- | --- | --- | --- | --- | --- | --- | --- |
|  | Mean | SE | RMSE | Type I | Mean | SE | RMSE | Coverage |
| Balanced Pleiotropy |  |  |  |  |  |  |  |  |
| IVW | -0.002 | 0.044 | 0.047 | 0.064 | 0.296 | 0.045 | 0.046 | 0.928 |
| MR-Egger | -0.005 | 0.130 | 0.137 | 0.068 | 0.253 | 0.132 | 0.142 | 0.918 |
| Median | -0.001 | 0.025 | 0.027 | 0.076 | 0.293 | 0.030 | 0.032 | 0.942 |
| Mode | 0.000 | 0.032 | 0.026 | 0.018 | 0.283 | 0.042 | 0.037 | 0.964 |
| Lasso | -0.001 | 0.016 | 0.022 | 0.146 | 0.294 | 0.019 | 0.026 | 0.850 |
| MR-Presso | -0.002 | 0.018 | 0.022 | 0.120 | 0.296 | 0.024 | 0.030 | 0.878 |
| MR-Raps | -0.002 | 0.041 | 0.042 | 0.052 | 0.302 | 0.043 | 0.041 | 0.954 |
| ConMix | -0.001 | — | 0.021 | 0.114 | 0.300 | — | 0.027 | 0.878 |
| JAM-MR | -0.001 | 0.021 | 0.024 | 0.092 | 0.291 | 0.025 | 0.033 | 0.874 |
| Oracle | -0.001 | 0.017 | 0.017 | 0.048 | 0.295 | 0.020 | 0.020 | 0.952 |
| Directional Pleiotropy, InSIDE Satisfied |  |  |  |  |  |  |  |  |
| IVW | 0.207 | 0.039 | 0.208 | 1.000 | 0.502 | 0.040 | 0.204 | 0.000 |
| MR-Egger | 0.005 | 0.112 | 0.114 | 0.052 | 0.253 | 0.116 | 0.131 | 0.918 |
| Median | 0.067 | 0.026 | 0.072 | 0.708 | 0.372 | 0.032 | 0.080 | 0.382 |
| Mode | 0.004 | 0.033 | 0.025 | 0.014 | 0.291 | 0.040 | 0.032 | 0.974 |
| Lasso | 0.044 | 0.017 | 0.050 | 0.678 | 0.358 | 0.020 | 0.066 | 0.250 |
| MR-Presso | 0.090 | 0.024 | 0.094 | 0.956 | 0.412 | 0.029 | 0.117 | 0.046 |
| MR-Raps | 0.173 | 0.039 | 0.175 | 1.000 | 0.478 | 0.041 | 0.180 | 0.000 |
| ConMix | 0.004 | — | 0.022 | 0.122 | 0.310 | — | 0.031 | 0.858 |
| JAM-MR | 0.015 | 0.021 | 0.034 | 0.166 | 0.320 | 0.027 | 0.051 | 0.808 |
| Oracle | 0.000 | 0.017 | 0.017 | 0.042 | 0.295 | 0.020 | 0.021 | 0.930 |
| Directional Pleiotropy, InSIDE Violated |  |  |  |  |  |  |  |  |
| IVW | 0.372 | 0.046 | 0.374 | 1.000 | 0.666 | 0.046 | 0.367 | 0.000 |
| MR-Egger | 0.687 | 0.124 | 0.699 | 1.000 | 0.946 | 0.127 | 0.660 | 0.002 |
| Median | 0.215 | 0.038 | 0.240 | 1.000 | 0.541 | 0.044 | 0.261 | 0.006 |
| Mode | 0.001 | 0.028 | 0.022 | 0.004 | 0.287 | 0.037 | 0.030 | 0.972 |
| Lasso | 0.068 | 0.019 | 0.076 | 0.840 | 0.397 | 0.022 | 0.106 | 0.074 |
| MR-Presso | 0.206 | 0.034 | 0.213 | 1.000 | 0.526 | 0.036 | 0.231 | 0.000 |
| MR-Raps | 0.342 | 0.049 | 0.344 | 1.000 | 0.642 | 0.049 | 0.344 | 0.000 |
| ConMix | 0.001 | — | 0.020 | 0.074 | 0.302 | — | 0.027 | 0.904 |
| JAM-MR | 0.027 | 0.022 | 0.123 | 0.096 | 0.358 | 0.028 | 0.189 | 0.822 |
| Oracle | 0.000 | 0.017 | 0.017 | 0.036 | 0.295 | 0.020 | 0.021 | 0.950 |

**Table 9:** Simulation results with sample sizes of *N*_1_ = *N*_2_ = 200000 and *P* = 200 genetic variants that explain 7% genetic variation in the risk factor.

| Method | Causal Effect $\theta = 0$ | | | | Causal Effect $\theta = 0.3$ | | | |
| --- | --- | --- | --- | --- | --- | --- | --- | --- |
|  | Mean | SE | RMSE | Type I | Mean | SE | RMSE | Coverage |
| Balanced Pleiotropy |  |  |  |  |  |  |  |  |
| IVW | 0.002 | 0.031 | 0.032 | 0.062 | 0.298 | 0.031 | 0.032 | 0.926 |
| MR-Egger | 0.007 | 0.092 | 0.093 | 0.062 | 0.261 | 0.093 | 0.104 | 0.916 |
| Median | 0.001 | 0.015 | 0.015 | 0.060 | 0.293 | 0.018 | 0.020 | 0.922 |
| Mode | 0.001 | 0.023 | 0.016 | 0.008 | 0.287 | 0.026 | 0.023 | 0.970 |
| Lasso | 0.001 | 0.010 | 0.010 | 0.064 | 0.296 | 0.011 | 0.015 | 0.872 |
| MR-Presso | 0.001 | 0.011 | 0.012 | 0.064 | 0.297 | 0.014 | 0.018 | 0.866 |
| MR-Raps | 0.002 | 0.028 | 0.027 | 0.040 | 0.301 | 0.029 | 0.028 | 0.954 |
| ConMix | 0.000 | — | 0.011 | 0.112 | 0.298 | — | 0.016 | 0.836 |
| JAM-MR | 0.000 | 0.011 | 0.012 | 0.066 | 0.294 | 0.014 | 0.018 | 0.858 |
| Oracle | 0.000 | 0.010 | 0.009 | 0.032 | 0.296 | 0.012 | 0.013 | 0.922 |
| Directional Pleiotropy, InSIDE Satisfied |  |  |  |  |  |  |  |  |
| IVW | 0.208 | 0.027 | 0.209 | 1.000 | 0.503 | 0.028 | 0.204 | 0.000 |
| MR-Egger | 0.003 | 0.079 | 0.083 | 0.058 | 0.261 | 0.080 | 0.093 | 0.912 |
| Median | 0.054 | 0.016 | 0.057 | 0.944 | 0.361 | 0.020 | 0.064 | 0.104 |
| Mode | 0.000 | 0.018 | 0.015 | 0.014 | 0.289 | 0.024 | 0.021 | 0.966 |
| Lasso | 0.029 | 0.010 | 0.032 | 0.742 | 0.342 | 0.012 | 0.046 | 0.142 |
| MR-Presso | 0.082 | 0.016 | 0.084 | 1.000 | 0.404 | 0.019 | 0.106 | 0.000 |
| MR-Raps | 0.171 | 0.027 | 0.172 | 1.000 | 0.474 | 0.028 | 0.175 | 0.000 |
| ConMix | 0.001 | — | 0.012 | 0.146 | 0.303 | — | 0.016 | 0.840 |
| JAM-MR | 0.005 | 0.011 | 0.013 | 0.088 | 0.303 | 0.014 | 0.022 | 0.888 |
| Oracle | -0.001 | 0.010 | 0.010 | 0.032 | 0.296 | 0.012 | 0.012 | 0.942 |
| Directional Pleiotropy, InSIDE Violated |  |  |  |  |  |  |  |  |
| IVW | 0.372 | 0.032 | 0.373 | 1.000 | 0.666 | 0.032 | 0.367 | 0.000 |
| MR-Egger | 0.705 | 0.086 | 0.710 | 1.000 | 0.974 | 0.088 | 0.679 | 0.000 |
| Median | 0.182 | 0.026 | 0.198 | 1.000 | 0.509 | 0.030 | 0.224 | 0.000 |
| Mode | 0.000 | 0.017 | 0.013 | 0.008 | 0.287 | 0.022 | 0.022 | 0.954 |
| Lasso | 0.050 | 0.011 | 0.053 | 0.950 | 0.370 | 0.013 | 0.074 | 0.020 |
| MR-Presso | 0.207 | 0.024 | 0.211 | 1.000 | 0.522 | 0.025 | 0.225 | 0.000 |
| MR-Raps | 0.340 | 0.034 | 0.341 | 1.000 | 0.639 | 0.034 | 0.340 | 0.000 |
| ConMix | 0.001 | — | 0.011 | 0.124 | 0.300 | — | 0.016 | 0.866 |
| JAM-MR | 0.002 | 0.011 | 0.011 | 0.046 | 0.297 | 0.014 | 0.036 | 0.918 |
| Oracle | 0.000 | 0.010 | 0.009 | 0.024 | 0.295 | 0.012 | 0.013 | 0.928 |

**Table 10:** Simulation results with the default settings and 20% of the genetic instruments being pleiotropic.

| Method | Causal Effect $\theta = 0$ | | | | Causal Effect $\theta = 0.3$ | | | |
| --- | --- | --- | --- | --- | --- | --- | --- | --- |
|  | Mean | SE | RMSE | Type I | Mean | SE | RMSE | Coverage |
| Balanced Pleiotropy |  |  |  |  |  |  |  |  |
| IVW | 0.001 | 0.018 | 0.018 | 0.058 | 0.295 | 0.018 | 0.019 | 0.928 |
| MR-Egger | 0.005 | 0.054 | 0.055 | 0.062 | 0.262 | 0.055 | 0.069 | 0.888 |
| Median | 0.000 | 0.009 | 0.009 | 0.036 | 0.293 | 0.011 | 0.013 | 0.914 |
| Mode | 0.000 | 0.025 | 0.013 | 0.006 | 0.288 | 0.022 | 0.021 | 0.954 |
| Lasso | 0.000 | 0.006 | 0.007 | 0.080 | 0.295 | 0.007 | 0.010 | 0.842 |
| MR-Presso | 0.000 | 0.007 | 0.007 | 0.062 | 0.295 | 0.008 | 0.011 | 0.860 |
| MR-Raps | 0.000 | 0.005 | 0.007 | 0.118 | 0.300 | 0.007 | 0.009 | 0.852 |
| ConMix | 0.000 | — | 0.008 | 0.242 | 0.299 | — | 0.010 | 0.770 |
| JAM-MR | 0.000 | 0.007 | 0.007 | 0.064 | 0.293 | 0.009 | 0.012 | 0.846 |
| Oracle | 0.000 | 0.006 | 0.006 | 0.054 | 0.296 | 0.008 | 0.009 | 0.884 |
| Directional Pleiotropy, InSIDE Satisfied |  |  |  |  |  |  |  |  |
| IVW | 0.140 | 0.017 | 0.140 | 1.000 | 0.437 | 0.017 | 0.137 | 0.000 |
| MR-Egger | 0.003 | 0.049 | 0.052 | 0.060 | 0.263 | 0.050 | 0.063 | 0.886 |
| Median | 0.030 | 0.010 | 0.031 | 0.894 | 0.330 | 0.012 | 0.032 | 0.270 |
| Mode | 0.000 | 0.016 | 0.012 | 0.006 | 0.289 | 0.020 | 0.018 | 0.952 |
| Lasso | 0.010 | 0.006 | 0.012 | 0.390 | 0.314 | 0.007 | 0.017 | 0.538 |
| MR-Presso | 0.028 | 0.008 | 0.029 | 0.942 | 0.341 | 0.010 | 0.042 | 0.028 |
| MR-Raps | 0.004 | 0.005 | 0.009 | 0.160 | 0.338 | 0.010 | 0.052 | 0.358 |
| ConMix | 0.001 | — | 0.008 | 0.198 | 0.302 | — | 0.010 | 0.778 |
| JAM-MR | 0.004 | 0.007 | 0.008 | 0.090 | 0.300 | 0.009 | 0.010 | 0.896 |
| Oracle | 0.000 | 0.006 | 0.006 | 0.030 | 0.297 | 0.008 | 0.009 | 0.918 |
| Directional Pleiotropy, InSIDE Violated |  |  |  |  |  |  |  |  |
| IVW | 0.271 | 0.021 | 0.271 | 1.000 | 0.567 | 0.021 | 0.268 | 0.000 |
| MR-Egger | 0.577 | 0.057 | 0.580 | 1.000 | 0.856 | 0.058 | 0.559 | 0.000 |
| Median | 0.065 | 0.011 | 0.066 | 1.000 | 0.375 | 0.013 | 0.076 | 0.002 |
| Mode | 0.000 | 0.016 | 0.012 | 0.012 | 0.288 | 0.020 | 0.020 | 0.956 |
| Lasso | 0.010 | 0.006 | 0.012 | 0.356 | 0.313 | 0.008 | 0.017 | 0.580 |
| MR-Presso | 0.062 | 0.009 | 0.063 | 1.000 | 0.371 | 0.012 | 0.073 | 0.002 |
| MR-Raps | 0.012 | 0.006 | 0.046 | 0.188 | 0.359 | 0.010 | 0.103 | 0.562 |
| ConMix | 0.000 | — | 0.008 | 0.180 | 0.300 | — | 0.010 | 0.818 |
| JAM-MR | 0.002 | 0.007 | 0.007 | 0.054 | 0.295 | 0.008 | 0.010 | 0.894 |
| Oracle | 0.000 | 0.006 | 0.006 | 0.030 | 0.295 | 0.008 | 0.009 | 0.892 |

**Table 11:** Simulation results with the default settings and 50% of the genetic instruments being pleiotropic.

| Method | Causal Effect $\theta = 0$ | | | | Causal Effect $\theta = 0.3$ | | | |
| --- | --- | --- | --- | --- | --- | --- | --- | --- |
|  | Mean | SE | RMSE | Type I | Mean | SE | RMSE | Coverage |
| Balanced Pleiotropy |  |  |  |  |  |  |  |  |
| IVW | 0.000 | 0.028 | 0.027 | 0.058 | 0.297 | 0.028 | 0.029 | 0.938 |
| MR-Egger | 0.008 | 0.082 | 0.085 | 0.058 | 0.261 | 0.083 | 0.090 | 0.930 |
| Median | 0.001 | 0.012 | 0.014 | 0.100 | 0.294 | 0.014 | 0.019 | 0.860 |
| Mode | 0.001 | 0.014 | 0.011 | 0.014 | 0.286 | 0.020 | 0.020 | 0.932 |
| Lasso | 0.001 | 0.008 | 0.010 | 0.122 | 0.296 | 0.009 | 0.013 | 0.832 |
| MR-Presso | 0.001 | 0.010 | 0.013 | 0.124 | 0.296 | 0.013 | 0.020 | 0.798 |
| MR-Raps | 0.001 | 0.008 | 0.016 | 0.340 | 0.304 | 0.015 | 0.024 | 0.734 |
| ConMix | 0.001 | — | 0.010 | 0.218 | 0.298 | — | 0.013 | 0.810 |
| JAM-MR | 0.001 | 0.010 | 0.011 | 0.112 | 0.294 | 0.012 | 0.017 | 0.846 |
| Oracle | 0.001 | 0.008 | 0.008 | 0.044 | 0.297 | 0.010 | 0.010 | 0.932 |
| Directional Pleiotropy, InSIDE Satisfied |  |  |  |  |  |  |  |  |
| IVW | 0.346 | 0.021 | 0.346 | 1.000 | 0.643 | 0.022 | 0.343 | 0.000 |
| MR-Egger | -0.001 | 0.061 | 0.064 | 0.072 | 0.269 | 0.062 | 0.072 | 0.902 |
| Median | 0.218 | 0.018 | 0.224 | 1.000 | 0.532 | 0.019 | 0.236 | 0.000 |
| Mode | 0.012 | 0.013 | 0.016 | 0.090 | 0.304 | 0.018 | 0.014 | 0.990 |
| Lasso | 0.230 | 0.013 | 0.233 | 1.000 | 0.552 | 0.013 | 0.255 | 0.000 |
| MR-Presso | 0.298 | 0.018 | 0.298 | 1.000 | 0.598 | 0.018 | 0.299 | 0.000 |
| MR-Raps | 0.339 | 0.023 | 0.339 | 1.000 | 0.640 | 0.024 | 0.341 | 0.000 |
| ConMix | 0.004 | — | 0.011 | 0.200 | 0.315 | — | 0.082 | 0.734 |
| JAM-MR | 0.007 | 0.010 | 0.013 | 0.142 | 0.307 | 0.013 | 0.034 | 0.862 |
| Oracle | 0.000 | 0.008 | 0.008 | 0.042 | 0.297 | 0.010 | 0.010 | 0.940 |
| Directional Pleiotropy, InSIDE Violated |  |  |  |  |  |  |  |  |
| IVW | 0.535 | 0.022 | 0.535 | 1.000 | 0.833 | 0.022 | 0.533 | 0.000 |
| MR-Egger | 0.794 | 0.062 | 0.796 | 1.000 | 1.071 | 0.064 | 0.774 | 0.000 |
| Median | 0.618 | 0.015 | 0.618 | 1.000 | 0.909 | 0.018 | 0.610 | 0.000 |
| Mode | 0.017 | 0.017 | 0.094 | 0.028 | 0.302 | 0.027 | 0.081 | 0.966 |
| Lasso | 0.581 | 0.010 | 0.582 | 1.000 | 0.870 | 0.011 | 0.571 | 0.000 |
| MR-Presso | 0.556 | 0.018 | 0.556 | 1.000 | 0.846 | 0.019 | 0.546 | 0.000 |
| MR-Raps | 0.542 | 0.024 | 0.542 | 1.000 | 0.842 | 0.025 | 0.542 | 0.000 |
| ConMix | 0.001 | — | 0.010 | 0.162 | 1.081 | — | 0.835 | 0.138 |
| JAM-MR | 0.686 | 0.018 | 0.694 | 0.996 | 0.961 | 0.020 | 0.669 | 0.000 |
| Oracle | 0.001 | 0.008 | 0.008 | 0.038 | 0.296 | 0.010 | 0.011 | 0.922 |

The simulations of Tables 7-9 illustrate that the performance of JAM-MR deteriorates when the GWAS sample sizes and proportion of variation in the risk factor are small. The algorithm performed rather poorly in the directional pleiotropy simulations of Table 7, where *N*_1_ = *N*_2_ = 50000 and only 3% variation in the risk factor was explained by the SNPs included in the analysis. Both the bias and the coverage properties of confidence intervals were affected. Some of the competing methods were subject to larger biases still, and the mode-based method was a clear winner in this scenario.

In the simulations of Table 8 (*N*_1_ = *N*_2_ = 100000 and 5% variation in the risk factor explained by the SNPs), the performance of JAM-MR improved and was close to that observed in the simulations in the main part of the paper, with slightly worse coverage properties in some settings. Further improvement was observed in the simulations of Table 9 (*N*_1_ = *N*_2_ = 200000 and 7% variation in the risk factor explained by the SNPs). These results seem to suggest that JAM-MR is more suitable for Mendelian randomization analyses of complex polygenic traits, for which large meta-GWAS have been conducted on hundreds of thousands of individuals and hundreds of associated genetic variants have been identified.

The results of simulations with varying numbers of pleiotropic instruments are reported in Tables 10-11. In the simulations of Table 10 with only 20% invalid instruments, there was an improvement in the performance of all methods. Estimates of the causal effect of interest were more accurate and Type I error rates were decreased as a result. The performance of JAM-MR improved in line with this trend: our algorithm was again one of the most accurate approaches for causal effect estimation, at the cost of a small amount of undercoverage for confidence intervals.

The simulation of Table 11 (50% invalid instruments) was quite challenging. JAM-MR was still able to offer accurate causal effect estimates in two out of the three scenarios, but failed to do so in the scenario where the InSIDE assumption was violated. We note that JAM-MR does not depend on the InSIDE assumption. The reason for its failure was that the SNP-confounder effects in our simulations were generated to be positive (*δ*_*j*_ ∼ *N* (0.4, 0.2^2^)). As a result, in the last simulation of Table 11, the pleiotropic variants were equal in number to the valid SNPs and were stronger instrument on average. In addition, univariate causal effect estimates for the pleiotropic variants were fairly homogeneous in that simulation. This led JAM-MR to identify the set of pleiotropic SNPs as “valid” and downweight the valid genetic variants instead. This is a difficult setting for any Mendelian randomization method. Indeed, the mode was the only approach that managed to identify the true causal effect in all simulation scenarios.

The performance of JAM-MR with smaller sample sizes and fewer genetic instruments (Table 7 was not very accurate. To explore the performance of the method further, we conducted an additional simulation. We used scenario 2 (directional pleiotropy, InSIDE satisfied) and generated three datasets with the following specifications:

- *N*_1_ = *N*_2_ = 50000 individuals in each GWAS and *P* = 50 genetic variants that explain 3% of the genetic variation in the risk factor. This is identical to scenario 2 of Table 7 hence the results reported are the same. It resembles a standard Mendelian randomization analysis for a complex trait, using GWAS studies with relatively small sample sizes.
- *N*_1_ = *N*_2_ = 50000 individuals in each GWAS and *P* = 50 genetic variants that explain 10% of the genetic variation in the risk factor. This could correspond to a Mendelian randomization analysis for a more specific risk factor, for which only a small number of genetic instruments with strong associations have been identified.
- *N*_1_ = *N*_2_ = 300000 individuals in each GWAS and *P* = 50 genetic variants that explain 3% of the genetic variation in the risk factor. This case might occur when studying a complex trait with relatively low genetic heritability.

The results of this simulation experiment are reported in Table 12. As in Table 7, the performance of JAM-MR was suboptimal in that scenario. On the other hand, increasing either the GWAS sample sizes or the proportion of genetic variation in the risk factor had a beneficial effect on the method’s performance, as JAM-MR was able to identify the true causal effect with decent accuracy and its Type I error rate inflation was rather small. This suggests that JAM-MR is only unreliable in applications where both the sample size and the proportion of genetic variation in the risk factor explained by the genetic variants are small. In practice it is often possible to incorporate genetic datasets from multiple sources into a Mendelian randomization analysis, by combining consortia meta-GWAS studies with available genetic biobanks, in order to increase sample sizes. We recommend that researchers do so when using the JAM-MR algorithm.

**Table 12:** Simulation results for scenario 2 (directional pleiotropy, InSIDE atisfied) with *P* = 50 SNPs and various values for the sample size and genetic variation in the risk factor.

| Method | Causal Effect $\theta = 0$ | | | | Causal Effect $\theta = 0.3$ | | | |
| --- | --- | --- | --- | --- | --- | --- | --- | --- |
|  | Mean | SE | RMSE | Type I | Mean | SE | RMSE | Coverage |
| $N_1 = N_2 = 50000, P = 50, GV = 3\%$ | | | | | | | | |
| IVW | 0.205 | 0.057 | 0.208 | 1.000 | 0.497 | 0.061 | 0.202 | 0.030 |
| MR-Egger | 0.004 | 0.161 | 0.169 | 0.066 | 0.220 | 0.168 | 0.194 | 0.916 |
| Median | 0.085 | 0.046 | 0.099 | 0.430 | 0.387 | 0.055 | 0.104 | 0.674 |
| Mode | 0.011 | 0.066 | 0.044 | 0.016 | 0.293 | 0.075 | 0.058 | 0.968 |
| Lasso | 0.074 | 0.029 | 0.086 | 0.654 | 0.389 | 0.034 | 0.103 | 0.344 |
| MR-Presso | 0.116 | 0.041 | 0.124 | 0.804 | 0.434 | 0.049 | 0.143 | 0.210 |
| MR-Raps | 0.177 | 0.059 | 0.181 | 0.958 | 0.485 | 0.063 | 0.190 | 0.082 |
| ConMix | 0.010 | — | 0.045 | 0.102 | 0.351 | — | 0.131 | 0.774 |
| JAM-MR | 0.064 | 0.069 | 0.120 | 0.270 | 0.368 | 0.062 | 0.110 | 0.684 |
| Oracle | -0.001 | 0.031 | 0.030 | 0.044 | 0.291 | 0.038 | 0.038 | 0.954 |
| $N_1 = N_2 = 50000, P = 50, GV = 10\%$ | | | | | | | | |
| IVW | 0.212 | 0.054 | 0.215 | 1.000 | 0.506 | 0.055 | 0.209 | 0.000 |
| MR-Egger | 0.010 | 0.164 | 0.172 | 0.058 | 0.267 | 0.164 | 0.180 | 0.916 |
| Median | 0.048 | 0.026 | 0.055 | 0.440 | 0.352 | 0.032 | 0.063 | 0.662 |
| Mode | 0.003 | 0.026 | 0.022 | 0.022 | 0.293 | 0.031 | 0.026 | 0.974 |
| Lasso | 0.023 | 0.016 | 0.031 | 0.320 | 0.328 | 0.019 | 0.040 | 0.670 |
| MR-Presso | 0.069 | 0.025 | 0.076 | 0.764 | 0.376 | 0.030 | 0.085 | 0.306 |
| MR-Raps | 0.173 | 0.053 | 0.176 | 0.990 | 0.472 | 0.055 | 0.176 | 0.016 |
| ConMix | 0.002 | — | 0.020 | 0.108 | 0.301 | — | 0.024 | 0.896 |
| JAM-MR | 0.005 | 0.017 | 0.021 | 0.106 | 0.303 | 0.021 | 0.033 | 0.882 |
| Oracle | 0.001 | 0.017 | 0.017 | 0.038 | 0.297 | 0.020 | 0.020 | 0.944 |
| $N_1 = N_2 = 300000, P = 50, GV = 3\%$ | | | | | | | | |
| IVW | 0.208 | 0.053 | 0.210 | 1.000 | 0.509 | 0.054 | 0.212 | 0.000 |
| MR-Egger | -0.009 | 0.163 | 0.181 | 0.080 | 0.286 | 0.167 | 0.189 | 0.912 |
| Median | 0.037 | 0.020 | 0.043 | 0.414 | 0.342 | 0.025 | 0.052 | 0.638 |
| Mode | 0.000 | 0.018 | 0.016 | 0.032 | 0.294 | 0.022 | 0.021 | 0.974 |
| Lasso | 0.013 | 0.013 | 0.021 | 0.208 | 0.315 | 0.015 | 0.026 | 0.738 |
| MR-Presso | 0.066 | 0.022 | 0.073 | 0.818 | 0.365 | 0.025 | 0.074 | 0.294 |
| MR-Raps | 0.166 | 0.052 | 0.170 | 0.982 | 0.469 | 0.053 | 0.173 | 0.014 |
| ConMix | 0.001 | — | 0.014 | 0.100 | 0.299 | — | 0.019 | 0.876 |
| JAM-MR | 0.001 | 0.013 | 0.014 | 0.088 | 0.298 | 0.015 | 0.019 | 0.886 |
| Oracle | 0.000 | 0.013 | 0.013 | 0.040 | 0.297 | 0.016 | 0.016 | 0.934 |

### Manhattan plots for the real-data application

Here we provide Manhattan plots obtained by running JAM-MR for the application in Section 4. Two Manhattan plots are given: one for systolic and one for diastolic blood pressure. In supplementary material, we provide the corresponding tables of results. These contain SNP names and chromosome positions, univariate summary statistics of association between SNPs and blood pressure traits, as well as coronary heart disease, and posterior inclusion probabilities assigned by JAM-MR to each SNP.

